# A Simple and Scalable Zebrafish Model of Sonic Hedgehog Medulloblastoma

**DOI:** 10.1101/2024.02.03.577834

**Authors:** Mattie J. Casey, Priya P. Chan, Qing Li, Cicely A. Jette, Missia Kohler, Benjamin R. Myers, Rodney A. Stewart

**Affiliations:** Department of Oncological Sciences, Huntsman Cancer Institute, University of Utah, Salt Lake City, UT 84112, USA; Department of Pediatrics, University of Utah School of Medicine, Salt Lake City, UT 84108, USA; Primary Children’s Hospital, Salt Lake City, UT 84113, USA; Department of Anatomic Pathology, University of Utah School of Medicine, Salt Lake City, UT 84112, USA; Department of Biochemistry, University of Utah School of Medicine, Salt Lake City, UT 84112, USA; Department of Bioengineering, University of Utah, Salt Lake City, UT 84112, USA

**Keywords:** Pediatric brain tumors, SHH medulloblastoma, zebrafish, PTCH1, GRK2, GRK3, midbrain-hindbrain boundary, TP53, CRISPR, valvula cerebelli

## Abstract

Medulloblastoma (MB) is the most common malignant brain tumor in children and is stratified into three major subgroups. The Sonic hedgehog (SHH) subgroup represents ∼30% of all MB cases and has significant survival disparity depending upon TP53 status. Here, we describe the first zebrafish model of SHH MB using CRISPR to mutate *ptch1*, the primary genetic driver in human SHH MB. These tumors rapidly arise adjacent to the valvula cerebelli and resemble human SHH MB by histology and comparative genomics. In addition, *ptch1-*deficient MB tumors with loss of *tp53* have aggressive tumor histology and significantly worse survival outcomes, comparable to human patients. The simplicity and scalability of the *ptch1* MB model makes it highly amenable to CRISPR-based genome editing screens to identify genes required for SHH MB tumor formation *in vivo*, and here we identify the *grk3* kinase as one such target.

## Introduction

Medulloblastoma (MB) is the most common malignant brain tumor to arise in children. It is frequently diagnosed between one and nine years of age but can occur in older children and even adults (Smoll & Drummond, 2012). MB arises in the cerebellum and often presents with symptoms of ataxia, compromised motor skills, and vision problems (Northcott et al., 2019). Depending on the presence of specific driver mutations and/or gene signatures within tumor cells at diagnosis, MB is classified into three main types: Wingless (WNT), Sonic hedgehog (SHH) and non-WNT/non-SHH (formerly known as Group 3 and Group 4). The latter two subgroups have poor overall survival (42-88%) compared to the WNT subgroup (97-100%) (Cavalli et al., 2017).

The most common genetic alteration in SHH MB is loss/mutation of *PTCH1* in ∼42% of cases (Skowron et al., 2021). SHH MB and WNT MB tumors often exhibit *TP53* mutations at diagnosis (Zhukova et al., 2013; Ramaswamy et al., 2017), and MB patients often relapse following the acquisition of mutations in *TP53* and amplification of *MYC/MYCN* from standard radiation and chemotherapy regimens (Hill et al., 2015). SHH MB patients, in particular, show a dramatic decrease in overall survival in response to *TP53* mutation (76-86% survival for *TP53*-wild-type and 32-50% survival for *TP53*-mutant) (Zhukova et al., 2013; Cavalli et al., 2017), thus prompting the World Health Organization to assign a more aggressive treatment regimen for these patients (Ramaswamy et al., 2017; Louis et al., 2021).

SHH MB has been well-characterized at the genomic level, with clear drivers of tumorigenesis arising from mutations that activate the SHH signaling pathway (Figure 1A). During normal embryonic development, the canonical SHH pathway is activated when the ligand SHH binds to the transmembrane receptor Patched 1 (PTCH1). PTCH1 is a negative regulator of the G protein-coupled receptor Smoothened (SMO). Thus, SHH indirectly activates SMO by inactivating PTCH1. Active SMO binds to and sequesters the catalytic subunit of protein kinase A (PKA), which normally inhibits SHH pathway signaling by phosphorylating and inactivating GLI transcription factors (Hammerschmidt et al., 1996; Niewiadomski et al., 2014; Arveseth et al., 2021; Happ et al., 2022; Walker et al., 2023). In the absence of SMO activity, suppressor of fused (SUFU) binds to GLI transcription factors and prevents their translocation to the nucleus. There are three GLI proteins: GLI1, which is a transcriptional activator that is only produced upon activation of the SHH pathway by GLI2 and GLI3, both of which act as transcriptional activators and repressors (Niewiadomski et al., 2019). During canonical SHH signaling, GLI2 activates and GLI3 represses SHH target genes. Therefore, when SMO is active, GLI is free to enter the nucleus and transactivate SHH pathway target genes that regulate CNS polarity and neural patterning (Hui & Angers 2011). In MB, inactivating mutations in *PTCH1* or *SUFU*, or activating mutations in *SMO* or *GLI2*, promote constitutive activation of the SHH pathway, leading to hyperplasia and tumorigenesis (Wang et al., 2022; Garcia-Lopez et al., 2021).

**Figure 1.**
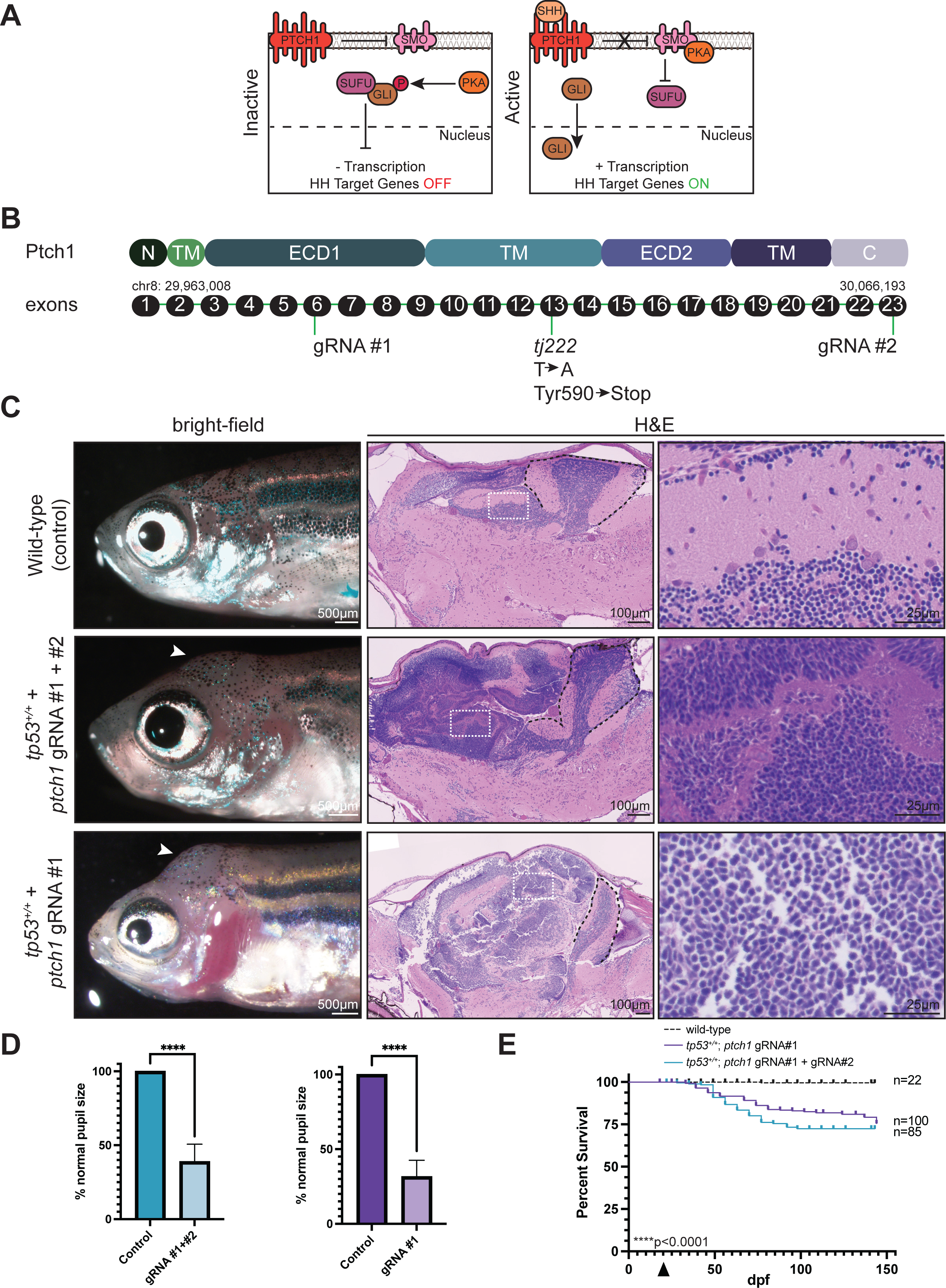
Transient *ptch1* crispants develop brain tumors. (A) Diagram of key mediators of the SHH pathway. (Left) Inactive SHH signaling: In the absence of SHH ligand, PTCH1 inhibits SMO. GLI is bound by SUFU and phosphorylated by PKA. GLI is unable to translocate to the nucleus, SHH target genes are not transcribed. (Right) Active SHH signaling: SHH ligand binds to PTCH1. SMO is activated, directly binds to PKA to prevent phosphorylation of GLI, and inhibits SUFU. GLI is free to translocate to the nucleus and transcribe SHH target genes. (B) Schematic of the Ptch1 protein domains and corresponding exons. gRNA target sites and the germline premature stop mutation (tj222) are indicated. The ptch1^tj222^ mutation (relevant to Figure 3) is also shown. The chromosomal location is noted above the first and last exon. N, N-terminal domain; TM, transmembrane domain; ECD1 and ECD2, extracellular domains 1 and 2; C, C-terminal domain. (C) (Left) Bright-field image of 6-wpf wild-type AB animals that were injected at the one-cell stage with the indicated *ptch1* gRNAs. Uninjected wild-type fish served as a negative control for this experiment. (Middle and right) Sagittal section of either control brain or *ptch1*-crispant brain tumors stained with hematoxylin and eosin. White arrowheads indicate the location of the tumor in the left panels. White boxes in the middle panel are shown at higher magnification in the right panel. Black dashed lines indicate the cerebellum. (D) Animals from the experiment described in C were analyzed for the presence of smaller pupils. The percentage of animals from each experimental group with normal pupil size was quantified. Data is plotted as the standard error of the mean. \*\*\*\**p*<0.0001. (E) Wild-type zebrafish were either left uninjected or were injected at the one-cell stage with either *ptch1* gRNAs #1 and #2 or *ptch1* gRNA #1 alone, and analyzed for survival following injection. Arrow indicates beginning of survival analysis.

Frontline treatment for children with MB includes tumor resection, radiation, and cytotoxic chemotherapy, which leads to devastating consequences for the developing brain (Robertson et al., 2006; Jakacki et al., 2012). For SHH MB, pharmacological inhibition of the SHH pathway, through targeting either SMO or downstream GLI transcription factors, has been challenging due to the development of drug resistance (Severini et al., 2020; Caimano et al., 2021). Drug resistance typically occurs through acquisition of secondary mutations in either SMO or downstream effectors, or an oncogenic program switch to alternate signaling pathways such as PI3K (Yauch et al., 2009; Kool et al., 2014; Dijkgraaf et al., 2011; Buonamici et al., 2010; Zhao et al., 2015). Moreover, many potential alternative therapies have limited efficacy due to their inability to cross the blood brain barrier (Terstappen et al., 2021). Current therapies therefore remain ineffective at improving long-term outcomes for patients, and the majority of *TP53*-mutant SHH MB patients eventually succumb to their disease.

Numerous SHH MB mouse models have been developed to better understand the etiology of SHH MB, to evaluate the response from targeted therapies and to identify additional signaling pathways for therapeutic exploitation (Roussel & Stripay 2020). Yet, current treatment regimens remain suboptimal. Efforts to improve treatment strategies and patient outcomes may require large-scale identification of new therapeutic targets and the analysis of combination therapies using a scalable animal model system like the zebrafish.

Here, we establish the first zebrafish model of SHH-pathway-driven brain tumors. Transient CRISPR/Cas9-mediated gene knockout of the *ptch1* gene in zebrafish leads to the rapid development of tumors that resemble SHH MB at the cellular and genomic levels. Similar to humans, combining *tp53* mutation with *ptch1* loss generates more aggressive tumors and reduced overall survival compared to *ptch1* loss alone. To test the potential for using multiplexed CRISPR (McCarty et al., 2020) in this model to identify new therapeutic targets for SHH MB, we injected zebrafish embryos with a combination of *ptch1* and *grk3* guide RNAs. Grk3 is the zebrafish homolog of GRK2, a downstream positive effector of the SHH pathway that has been previously reported to promote MB growth (Pusapati et al., 2018; Pathania et al., 2019). Indeed, loss of *grk3* significantly improved the survival of *ptch1*-crispant animals, indicating that Grk3 may represent a promising new therapeutic target for SHH MB. Moreover, our data provides a proof of principle showing that multiplexed CRISPR approaches could be combined with this highly scalable and low-cost SHH MB zebrafish model for the rapid identification of new therapeutic targets and combination treatments for the most aggressive MB subgroups.

## Results

### Transient *ptch1*-crispant zebrafish develop brain tumors

To analyze the consequence of *ptch1* loss in zebrafish, we knocked out *ptch1* using CRISPR/Cas9-mediated gene editing. The zebrafish Ptch1 protein shows 73% identity with human PTCH1 and is composed of a sterol-sensing domain in its transmembrane region, and two extracellular domains required for ligand recognition (Figure 1B, Zhang & Beachy 2023). We therefore designed guide RNAs (gRNAs) to target the first extracellular domain and the C-terminal domain of the *ptch1* gene (gRNAs #1 and #2, respectively, see Figure 1B). We injected one or both *ptch1* gRNAs into one-cell-stage wild-type AB embryos and subsequently monitored them for tumor formation. Brain protrusions were evident by 3-4 weeks post fertilization (wpf) in fish with *ptch1* single and double-injected gRNA’s (called crispants), with ∼76% of injected animals exhibiting obvious tumors and abnormal swimming behavior (data not shown) by 150 days (Figure 1C). Hematoxylin and eosin staining of the brain protrusions revealed cells with high nuclear-to-cytoplasmic ratios, and scant to near-absent cytoplasm and hyperchromatic nuclei suggestive of an embryonal neoplasm (or round blue cell tumor) (Figure 1C). Smaller pupil sizes were also observed in at least one eye in 25 out of 38 tumor-bearing, 5 to 8-week-old, *ptch1*-crispant animals, consistent with previous reports showing *ptch1* is required for optic vesicle development (Figure 1D, Bibliowicz & Gross 2009; Koudijs et al., 2005). Survival was decreased in double *ptch1-*crispant animals compared to single *ptch1* crispants (Figure 1E). We confirmed that the *ptch1* gene was disrupted in the *ptch1*-crispant tumors via sequencing (Figure S1A).

### Tp53 loss promotes aggressiveness in transient ptch1-crispant brain tumors in zebrafish

In murine models with either germline *Ptch1* mutation or somatic *Ptch1* disruption, *Tp53* loss accelerates tumor incidence and onset (Wetmore et al., 2001; Zuckermann et al., 2015). To determine whether Tp53 functions as a tumor suppressor in the zebrafish *ptch1* crispants, we injected one or both *ptch1* gRNAs into one-cell-stage embryos from a cross between *tp53*-mutant heterozygotes (*tp53^M214K^*, Berghmans et al., 2005) (Figure 2A) and subsequently monitored them for tumor formation. Survival was decreased in both single and double *ptch1*-crispant animals compared to control animals (*tp53^M214K^*;Casper), and homozygous mutations in *tp53* further exacerbated this phenotype (Figure 2B-C), mimicking the decreased survival outcomes in human MB patients (Zhukova et al., 2013; Cavalli et al., 2017) and mammalian model systems (Wetmore et al., 2001; Zuckermann et al., 2015). The mutant-*tp53*-mediated decrease in survival also correlated with an aggressive histopathology, as the tumors contained larger neoplastic cells with increased nuclear molding and the formation of discrete tumor nodules (Figure 2D, Figure S1B-C).

**Figure 2.**
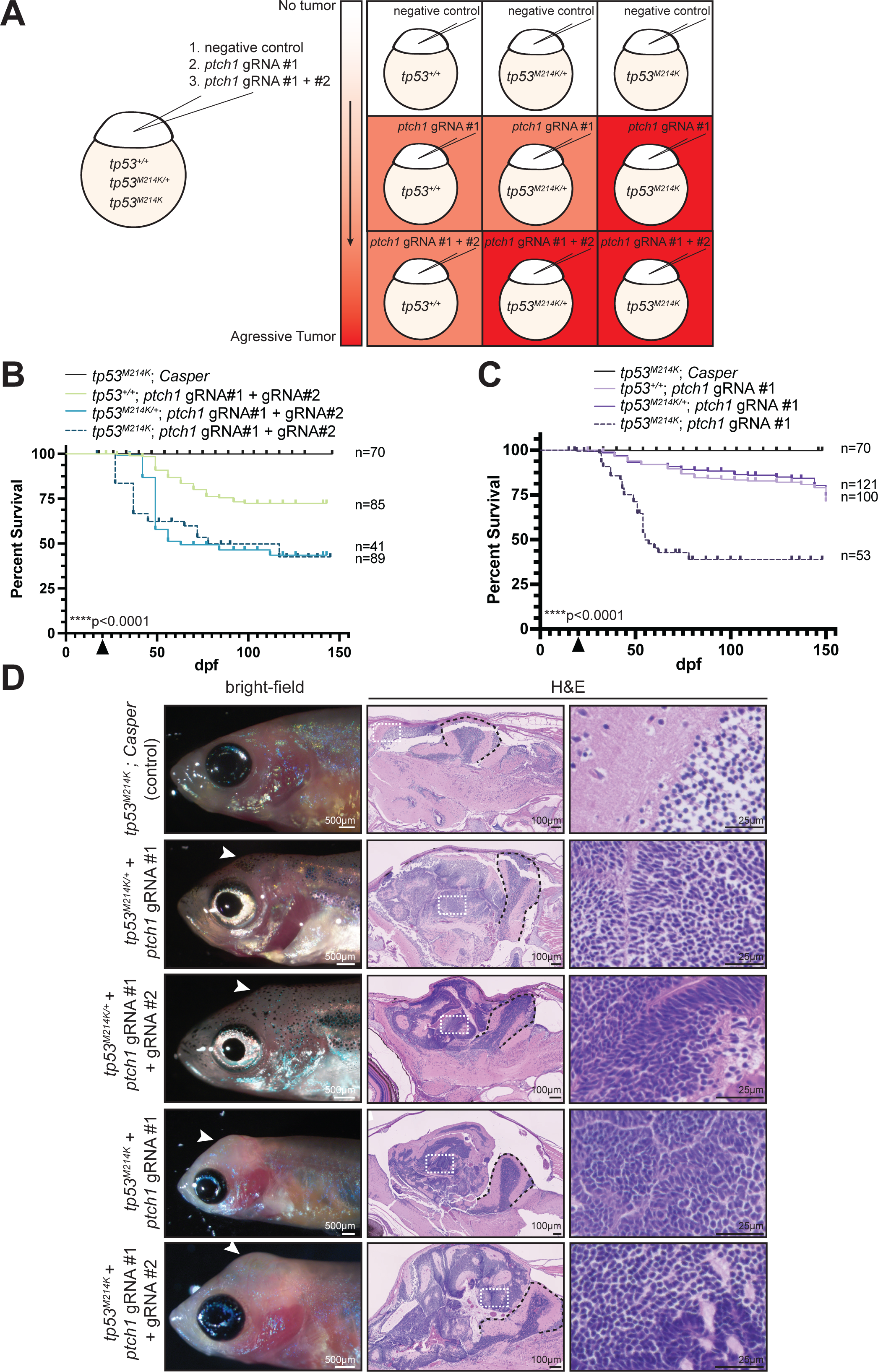
Mutation in *tp53* promotes aggressiveness in *ptch1*-crispant brain tumors. (A) Schematic of the embryo injection strategy to determine whether *tp53* acts as a tumor suppressor in the *ptch1*-crispant brain tumors. (B-C) Zebrafish from the indicated genetic backgrounds were injected at the one-cell stage with either *ptch1* gRNAs #1 and #2 (B) or *ptch1* gRNA #1 alone (C), and analyzed for survival following injection. Arrow indicates beginning of survival analysis. (D) (Left) Bright-field image of 7-wpf animals with the indicated genetic backgrounds that were injected at the one-cell stage with either *ptch1* gRNA #1 alone or *ptch1* gRNAs #1 and #2. (Middle and right) Sagittal section of control brain or *ptch1*-crispant brain tumors stained with hematoxylin and eosin. White arrowheads indicate location of the tumor in the left panels. White boxes in the middle panel are shown at higher magnification in the right panel. Black dashed lines indicate the cerebellum.

### Adult germline ptch1 mutants develop brain tumors

To verify that loss of *ptch1* causes brain tumors in zebrafish, we next tested whether germline loss of *ptch1* also gives rise to brain tumors. Due to the short lifespan and compromised health of the transient *ptch1-*crispant adult animals (Figure 1), it was not possible to establish a stable line using our *ptch1*-crispant methodology. We therefore obtained and analyzed a previously established *ptch1^tj222^* line (Figures 1A and 3) that has primarily been studied during embryonic development (Heisenberg et al., 1996; Koudijs et al., 2008). As expected from previous reports, most of the *ptch1^tj222^* homozygotes died at early developmental stages (data not shown); however, we identified four homozygous mutant escapers out of 147 adults (∼37 homozygotes expected from normal Mendelian ratios) from an incross between *ptch1^tj222^* heterozygotes. All four homozygous mutants developed brain tumors that resemble *ptch1*-crispant tumors, and tumorigenesis was independent of *tp53* status (Figure 3). This independent analysis using a stable *ptch1*-null line supports the conclusion that loss of *ptch1* drives the development of brain tumors in zebrafish.

**Figure 3.**
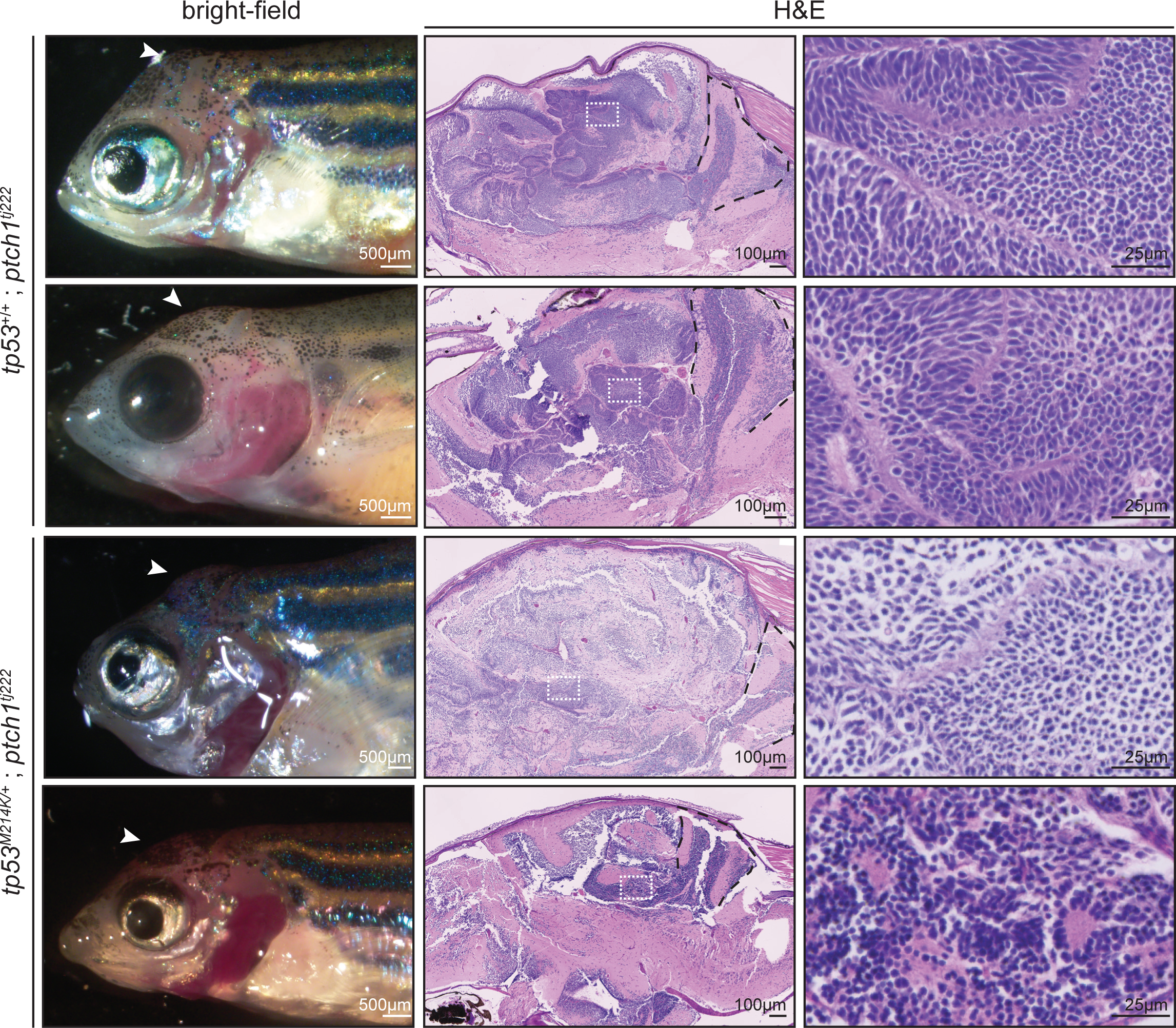
Zebrafish germline *ptch1*-mutant adults develop brain tumors. (Left) Bright-field image of the germline *ptch1^tj222^* mutant animals that survived to adulthood. (Middle and right) Sagittal section of *ptch1^tj222^*brain tumors stained with hematoxylin and eosin. White arrowheads indicate location of the tumor in the left panels. White boxes in the middle panel are shown at higher magnification in the right panel. Black dashed lines indicate the cerebellum.

### Ptch2 loss does not induce brain tumors in zebrafish

*PTCH2* mutations have also been reported in a small fraction of MB patients (Smyth et al., 1999). To determine if loss of *ptch2* induces brain tumors in zebrafish, we designed a gRNA targeting the second extracellular domain of *ptch2* and identified two independent mutations that are predicted to truncate the Ptch2 protein (Figure S2A-B). Transient *ptch2* crispants grew to fertile adult zebrafish with no evidence of tumor formation (data not shown) but with an obvious coloboma phenotype (Figure S2D). Stable *ptch2* crispant lines were isolated and crossed into a *tp53*-mutant background to maximize the potential for tumor formation in response to *ptch2* loss. Each *ptch2*-heterozygous, *tp53*-homozygous line was incrossed to generate 25% double-homozygous-mutant embryos for analysis. As previously reported for homozygous loss of *ptch2*, *ptch2/tp53*-double-homozygotes were largely embryonic lethal and also exhibited a coloboma phenotype (Figure S2C-D, Koudijs et al., 2008; Gordon et al., 2018). However, five percent of the double-homozygous-mutant animals escaped embryonic lethality and grew to adulthood. No tumors were observed among both heterozygous and escaper-homozygous *ptch2*-mutant adults in the *tp53*-mutant background. Thus, *ptch2* loss does not appear to induce brain tumors in zebrafish.

### Ptch1-mutant tumors genetically resemble human SHH MB

To determine the degree to which the zebrafish *ptch1-*crispant brain tumors resemble human embryonal tumors, we performed RNA sequencing to compare normal zebrafish brain tissue to the *ptch1-*mutant tumors (Figure 4A). The brains of tumor-bearing *ptch1*; *tp53^M214K^* animals were dissected and compared to *tp53^M214K^*; Casper control brains. Genes that were differentially expressed between the two tissues were analyzed by gene set enrichment analysis (GSEA). GSEA indicated a small number of enriched pathways, the most significant of which included genes either expressed in pediatric cancer or regulated by the SHH pathway (Figure 4B). Since *PTCH1* mutations give rise to SHH MB in humans, we compared the transcriptional signatures of zebrafish *ptch1*-crispant tumors to published human MB signatures using principal-component analysis (PCA) (GSE85217; Cavalli et al., 2017) and found that the zebrafish brain tumors tightly cluster with human SHH MB signatures, but not other MB subtype signatures (Figure 4C), showing *ptch1-*deficient zebrafish tumors most closely resemble SHH MB at the genomic level.

**Figure 4.**
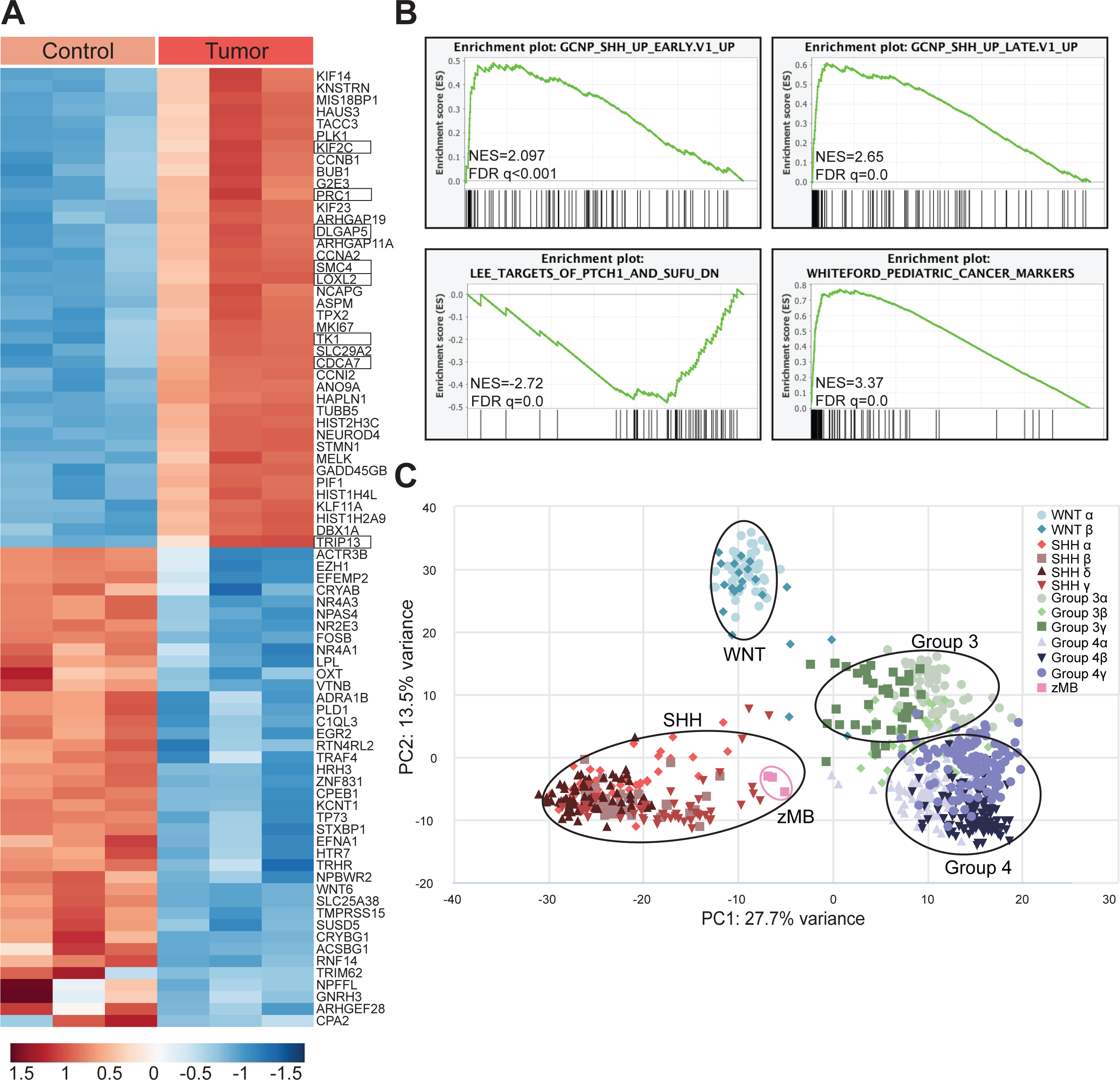
Zebrafish *ptch1*-crispant brain tumors resemble human SHH medulloblastoma. (A) Heatmap of the top 40 differentially regulated genes in *tp53^M214K^* control brain tissue (“Control”) and *tp53^M214K^*; *ptch1* gRNA #1 brain-tumor tissue (“Tumor”). Three biological replicates were analyzed per group. SHH pathway response genes are boxed. (Zhao et al. 2002 PNAS PMID 11960025). (B) Gene set enrichment analysis (GSEA) was performed to identify expression pathways enriched in *tp53^M214K^*; *ptch1-*crispant, brain-tumor tissue compared to *tp53^M214K^* control brain. NES, normalized enrichment score; FDR, false discovery rate. (C) Principal component analysis comparing zebrafish *tp53^M214K^*; *ptch1-*crispant brain tumors and human MB samples (Cavalli et al., 2017).

To identify potential cellular origins for the *ptch1*-mutant brain tumors, we compared genes differentially expressed in our *ptch1*-mutant brain tumors to that of our previously published CNS NB-FOXR2 pediatric brain tumor model (Modzelewska et al., 2016). The CNS NB-FOXR2 brain tumor model originates from the expression of either wild-type or oncogenic NRAS in oligodendrocyte precursor cells (OPCs). As expected, the CNS NB-FOXR2 brain tumors revealed significant upregulation of OPC markers *sox10* and *erbb3* (Figure S3A, GSE80768; Modzelewska et al., 2016). In contrast, *ptch1*-mutant brain tumors were positively enriched for ciliogenesis and neuronal, specifically glutamatergic, markers and lacked expression of oligodendrocyte markers (Figure S3B), indicating that the *ptch1*-crispant brain tumors have characteristics of neural progenitor/stem cells and likely arise from a different cell of origin than CNS NB-FOXR2 tumors. These data are consistent with the hypothesis that *ptch1*-mutant brain tumors originate in a neural stem cell compartment in the cerebellum, similar to human SHH MB.

### Zebrafish ptch1-mutant tumors arise near the midbrain-hindbrain boundary

A morphological feature of the *ptch1*-crispant tumors is a “bump” on the head, usually located anterior to the cerebellum (Figure 1). Histologically, we also observe aggressive infiltration of *ptch1*-crispant tumor cells coming from the anterior cerebellar region toward the 3^rd^ ventricular space at the mid-hindbrain boundary (MHB) region (Figure 1). In addition, the structure of the corpus cerebelli and lobus caudalis cerebelli (located in the posterior region of the hindbrain, Figure S1B) appeared relatively normal in the majority of *ptch1*-crispant animals (see Figure 1C and Figure 2D). Based on these anatomical observations, we hypothesized that the *ptch1*-crispant tumors arise in the valvula cerebelli, a unique anterior extension/lobe of the teleost cerebellum adjacent to midbrain, or the MHB subventricular region, or both. To test this, we analyzed candidate regions of the zebrafish cerebellum that are responsive to hedgehog signaling and therefore might contain the cell of origin for the *ptch1*-crispant brain tumors, by using available transgenic lines with green fluorescent SHH-producing cells (*Tg*(*shh:H2A-GFP*)) and red fluorescent SHH-responsive cells (*Tg*(*8xGli:mCH*)) (Gordon et al., 2018; Mich et al., 2014). We generated double-transgenic lines to visualize these two cell types within the zebrafish MHB and cerebellum (Figure S4A-B). Interestingly, we found that SHH-producing and - responding cells were tightly juxtaposed at the MHB-anterior cerebellar region (Figure S4A-B). Considering that cerebellar granule neural progenitor (GNP) cells are the cell of origin in human SHH MB, and GNP cells arise from the valvula cerebelli in zebrafish, we reasoned that the valvula cerebelli contains the cell of origin for the *ptch1*-crispant tumors. To evaluate this model, we compiled a list of genes that are known to be expressed in the zebrafish MHB region (Table S3) and analyzed whether they are enriched in *ptch1*-crispant brains compared to control brains. Indeed, this analysis revealed positive enrichment for anterior cerebellar/MHB-expressing genes within the *ptch1*-crispant brains (Figure S5A-B) suggesting that the brain tumors we observe in *ptch1*-mutant zebrafish likely originates in the region of the valvula cerebelli (see Discussion).

### SHH MB tumorigenesis requires Grk3

With the ultimate goal of using the *ptch1* crispants for target discovery in high throughput screens, we investigated whether CRISPR-based genome editing could be a useful tool to rapidly identify important mediators of SHH MB in fish (McCarty et al., 2020). Since SMO is downstream of PTCH1 in the SHH pathway, we designed a gRNA to knock out the *smo* gene as a positive control. The survival of zebrafish injected with *ptch1* gRNA alone, or the combination of *ptch1* and *smo* gRNAs, was monitored over a 60-day period (Figure 5A). As expected, loss of *smo* in *ptch1* crispants significantly improved survival compared to control gRNAs, albeit incompletely which is likely due to incomplete CRISPR cutting and early embryonic requirements for *smo* in tissue patterning (Figure 5A).

**Figure 5.**
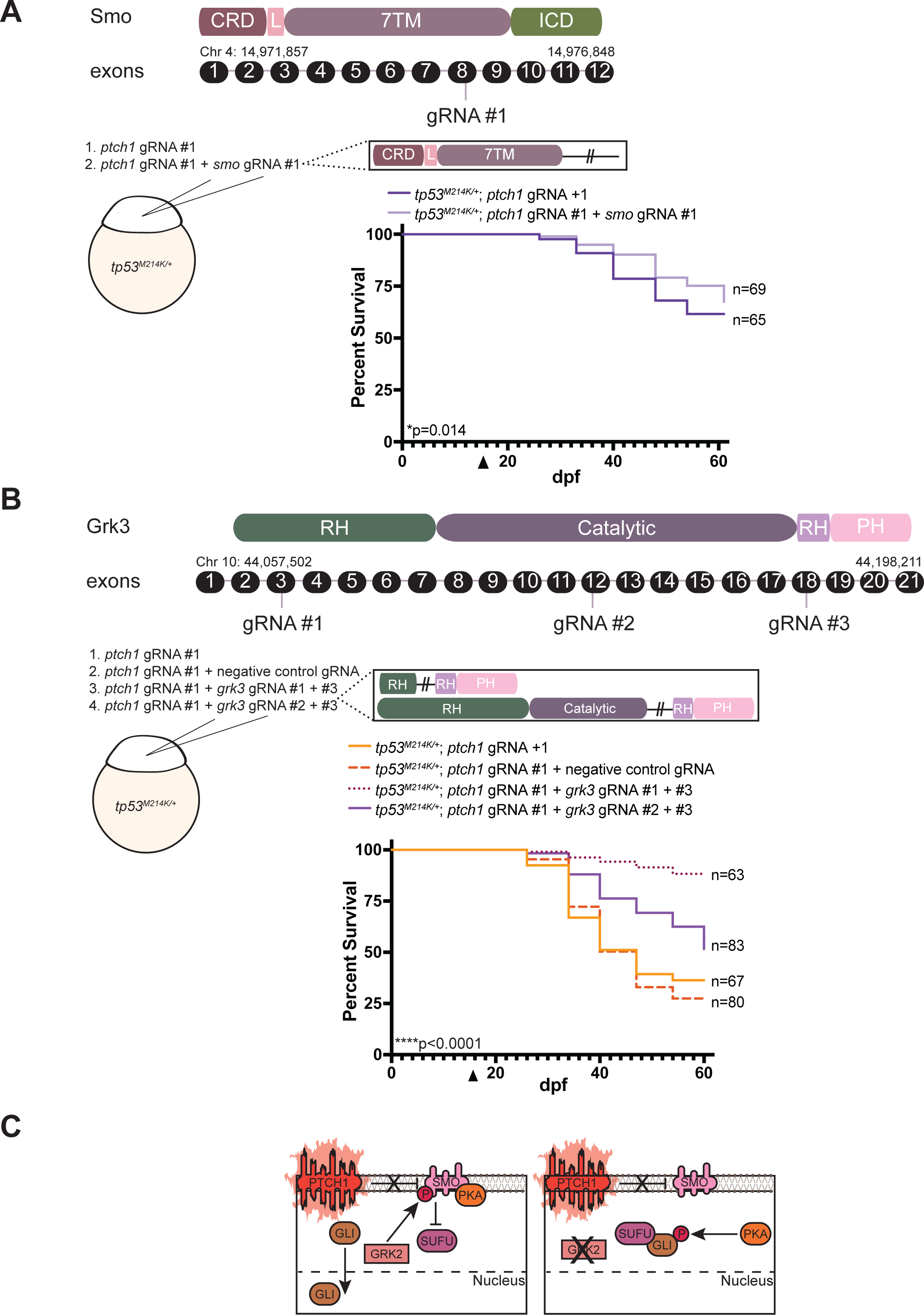
Loss of *grk3* improves overall survival of *ptch1* animals. (A) (Top) Schematic of the Smo protein domains and corresponding exons. The gRNA target site is indicated. CRD, cysteine-rich domain; L, linker domain; 7TM, seven transmembrane domain; ICD, intracellular domain. (Middle) Schematic of injection strategy and predicted zebrafish Smo mutant protein. (Bottom) Survival curve of *ptch1* crispants compared to *ptch1/smo* double crispants. Arrow indicates beginning of survival analysis. (B) (Top) Schematic of the Grk3 protein domains and corresponding exons. gRNA target sites are indicated. RH, RGS-homology domain; Catalytic, catalytic domain; PH, pleckstrin-homology domain. (Middle) Schematic of injection strategy and predicted zebrafish Grk3 mutant protein. (Bottom) Survival curve of *ptch1* crispants and *ptch1/*negative control crispants compared to *ptch1/grk3* double crispants using two independent *grk3* gRNAs. Arrow indicates beginning of survival analysis. (C) Schematic describing how loss of GRK3 disrupts SHH-induced tumorigenesis. (Left) In *ptch1*-deficient MB tumors, SMO is not inhibited by mutant PTCH1 promoting phosphorylation by GRK, which enables SMO to bind and sequester PKA. In turn, GLI is no longer modulated by PKA-mediated phosphorylation or sequestered by SUFU and is instead translocated to the nucleus to constitutively transcribe SHH target genes. (Right) Loss/inhibition of GRK2 prevents SMO phosphorylation, thereby preventing SMO from sequestering PKA. Free PKA phosphorylates GLI which now binds to SUFU and is unable to translocate to the nucleus. SHH target genes are no longer transcribed.

We next identified candidates for new SHH-MB therapeutic targets. We prioritized the analysis of GRK2, a G protein-coupled receptor kinase that promotes activation of the SHH signaling pathway through phosphorylation of SMO (Chen et al., 2011; Arveseth et al., 2021; Walker et al., 2023). GRK2 has been shown to stimulate the growth and proliferation of MB cell lines (Pathania et al., 2019; Pusapati et al., 2018) but has never been tested *in vivo* for its importance in SHH-pathway-induced tumorigenesis. To knock out the zebrafish *GRK2* homolog, called *grk3,* we designed gRNAs to target the catalytic and RGS-homology (RH) domains of *grk3* (Figure 5B). We injected zebrafish one-cell-stage embryos with the *ptch1* gRNA either alone or in combination with *grk3* gRNAs, and subsequently monitored their survival (Figure 5B). Similar to loss of *smo,* we found that loss of *grk3,* via two independent *grk3* gRNAs, significantly improved the lifespan of *ptch1*-crispant animals (Figure 5B) compared to animals injected with *ptch1* plus negative-control gRNAs (Figure 5B). These findings highlight the potential future utility of a transient, multiplexed-CRISPR-based approach to generate large numbers of *ptch1*-mutant, brain-tumor-bearing animals for drug discovery.

## Discussion

Pre-clinical models of SHH-driven MB are an important tool for identifying and testing new strategies to eliminate high risk tumors. Several robust mouse models of SHH-driven MB, including the high-risk *TP53* subgroup, have been used to enhance our understanding of the genetic and epigenetic drivers underlying tumor formation as well as for pre-clinical testing of SHH pathway inhibitors, such as vismodegib and sonidegib (reviewed in Roussel & Stripay 2020). While murine models will continue to be valuable in identifying and testing rationally designed treatments, these models are not ideal for high-throughput screening approaches, such as genetic and drug screens, due to small litter sizes and high costs associated with scalability. Unbiased screening approaches in animals are needed to complement mechanism-based approaches to identify both tumor- and host-dependent vulnerabilities that overcome the shortcomings of the current generation of SHH pathway inhibitors, such as rapid drug resistance and developmental toxicities. As a first step to addressing this problem, we have established a simple and scalable Crisper/Cas9-based method to generate *ptch1*-deficient zebrafish models of SHH-driven MB that recapitulate the histology and genomic signatures of human SHH MB.

Previous studies of *ptch1* in zebrafish have focused on its requirement during organogenesis of the eye, ear and fin (Heisenberg et al., 1996; Whitfield et al., 1996; van Eeden et al., 1996; Chassaing et al., 2016). In addition, *ptch1 loss* was shown to modify the malignancy of *notch1-*driven T-cell acute lymphoblastic leukemia in zebrafish tumor models (Burns et al., 2018). Here, we demonstrate that germ-line or somatic loss of *ptch1* alone is sufficient to drive formation of embryonal brain tumors in zebrafish that resemble SHH MB at both histological and genomic levels, thus demonstrating that *ptch1* is a highly conserved tumor suppressor in brain tumor etiology. In addition, we show that the combined loss of *ptch1* and *tp53* significantly enhances tumor malignancy and decreases survival, consistent with observations in murine MB models as well as and human SHH MB patients (Ramaswamy et al., 2017; Zhukova et al., 2013; Wetmore et al., 2001). These data also suggest that zebrafish may succumb to other SHH-driven tumors, such as Rhabdomyosarcoma and basal cell carcinoma (Pietrobono et al., 2019), which we did not detect in this study due to the severity and early lethality of the brain tumors, but could be tested in the future using conditionally targeted *ptch1* alleles.

Granule neural precursor cells (GNPs) are an established cell of origin for SHH-driven MB in mammals (Schuller et al., 2008; Yang et al., 2008; Kani et al., 2010). During cerebellar development, mammalian GNPs arise from the rhombic lip and migrate tangentially to cover the surface of the dorsal cerebellum to form the external granule layer (EGL) (Machold & Fishell 2005; Wang et al., 2005). SHH secretion from underlying Purkinje cells induces proliferation of the EGL that occurs from embryogenesis until puberty, which is required to produce enough GNPs to sustain the growth of the cerebellum during development and attain its final size for the rest of adult life. Sustained SHH signaling in GNPs (for example, through *PTCH1* loss) maintains a proliferative state in the EGL and prevents differentiation, thereby promoting MB formation in infants and adolescents. In contrast, zebrafish lack the transient EGL proliferation zone during embryogenesis and Shh pathway components (e.g., *shh, gli1/2/3, ptch1*) are absent from the dorsal cerebellum (Chaplin et al., 2010; Schwend et al., 2010; McFarland et al., 2008). Thus, the formation of MB in zebrafish is somewhat surprising given the differences in neuroanatomy and suggests that zebrafish MB arises from an anatomical location that is distinct from mammals. Nevertheless, because these zebrafish brain tumors involve the same molecular (SHH pathway activation) and cellular (EGL/GNP proliferation) mechanisms as their mammalian counterparts, we expect them to be broadly useful for the study of MB.

We hypothesize that zebrafish SHH-driven MB arises in the valvula cerebelli, a unique anterior lobe of the teleost cerebellum that is an extension of the corpus cerebelli located underneath the optic tectum adjacent to the midbrain ventricle cavity (Schuller et al., 2008; Yang et al., 2008; Kani et al., 2010). The valvula cerebelli maintains the same layered structure as the corpus cerebelli (molecular, Purkinje and granule-neuron layers) and produces *atoh1a*-positive GNPs that migrate into the granule neuron layer throughout the life of the fish (Consalez et al., 2021; Bae et al., 2009; Kaslin et al., 2009; Hashimoto & Hibi 2012; Hibi et al., 2017). Indeed, a unique feature of the zebrafish cerebellum is its ability to regenerate and grow throughout adulthood due to the presence of cerebellar stem-cell niches that continually generate GNPs, so a transient period of EGL proliferation is not necessary during embryogenesis (Consalez et al., 2021; Bae et al., 2009; Kaslin et al., 2009; Hashimoto & Hibi 2012; Hibi et al., 2017). In addition, the anterior-ventral location of the valvula cerebelli places it in proximity of Shh-producing cells in the ventral neural tube (Schuller et al., 2008; Yang et al., 2008; Kani et al., 2010), and we show that zebrafish express both Shh-producing and Shh-responsive cells at the border of the cerebellum and MHB. The location of the zebrafish MB tumors by gross morphology and histology shows a preponderance of tumors arising at the midbrain boundary region and expanding via the mesencephalic cavity to dorsal regions. Finally, our genomic analysis shows transcriptional signatures consistent with SHH-induced GNP proliferation as well as cerebellar and MHB cell types. Thus, we propose that the continuous supply of GNPs in the valvula cerebelli are an origin for MB tumor formation in zebrafish, which could be tested in the future using conditional systems to inactivate *ptch1* in different cerebellar compartments.

In this study, we have outlined methods to establish SHH MB tumors in zebrafish which develop obvious masses within 3-4 weeks and can be scaled to hundreds of animals each day by a single user. For rapid identification of genes required for tumor formation and survival, we utilized a dual CRISPR-based approach to genetically modify potential therapeutic targets. These experiments support the idea that multiple gRNAs can be injected into embryos with the *ptch1* gRNAs to perform an unbiased screen for genes that impact tumor growth while at the same time assessing embryonic and developmental toxicities that currently hamper the use of single-agent SHH inhibitors. Here we applied this approach to study the role of GRK2 (zebrafish Grk3) in MB tumorigenesis, which we recently demonstrated plays a role in SHH signaling by phosphorylating SMO to promote PKA recruitment and inactivation which triggers GLI activation (Figure 5C; Walker et al., 2023; Arveseth et al., 2021; Happ et al., 2022). Loss of *grk3* in zebrafish causes stereotypical *shh*-deficient developmental phenotypes, such as cyclopia (Zhao et al., 2016). However, the potential role of *grk3* in SHH MB tumorigenesis *in vivo* is not known. Here, we show CRISPR-mediated inactivation of *grk3* significantly improved survival of *ptch1* crispants, demonstrating that Grk3 represents a key mediator of oncogenic SHH signaling *in vivo* as well as a potential therapeutic target (Pusapati et al., 2018). Based on our findings we propose that GRK2 small molecule inhibitors, which include Cmpd101, 14AS, and paroxetine (an FDA-approved selective serotonin re-uptake inhibitor) and are currently used in cardiovascular disease models, should be evaluated in conventional murine models of SHH-driven MB (Schumacher et al., 2015; Ikeda et al., 2007; Thal et al, 2012; Waldschmidt et al., 2017).

## Acknowledgements

We thank the Bruce Appel, Richard Dorsky and Kristen Kwan labs for reagents, and Kazuyuki Hoshijima, Annika Thorpe, and Mya Scheib for technical support. We thank Christian Davidson for initial pathology consultation. Research reported in this publication utilized the High-Throughput Genomics, Bioinformatic Analysis, and Biorepository and Molecular Pathology Shared Resources at the University of Utah Huntsman Cancer Institute, which is supported by the National Cancer Institute (P30CA042014). The computational resources used were partially funded by the NIH Shared Instrumentation Grant 1S10OD021644-01A1. We acknowledge the Cell Imaging Core at the University of Utah for use of the Zeiss Axio Scan.Z1 slide scanner and thank Xiang Wang for assistance in image acquisition. We thank current and former members of the RAS laboratory for thoughtful discussions, and the zebrafish core at the University of Utah for providing animal husbandry. This work was supported by funding from the National Institutes of Health (R01NS106527, P30CA042014) (RAS), as well as the Huntsman Cancer Foundation (RAS) and American Cancer Society (RSG-22-077-01-CCB) (BRM).

## Author contributions

This study was conceived and designed by MJC and RAS. MJC and PPC performed the experiments. MJC, QL, MK and RAS analyzed the data. The manuscript was prepared by MJC, CAJ, BRM, and RAS with input from all authors.

## STAR Methods

Resource Availability

### Lead contact

Further information and requests for resources and reagents should be directed to Rodney Stewart.

### Materials availability

This study did not generate new unique reagents.

### Data and code availability

- Raw RNA sequencing data is publicly available and has been deposited at GEO under the acquisition number GSE242897. Analyzed RNA sequencing data is available via supplemental Table S1.
- This paper does not report any original code.
- Any additional information required to reanalyze the data reported in this paper is available from the lead contact upon request.

## Experimental Model and Subject Details

### Ethics statement

All experiments using zebrafish conformed to the regulatory standards and guidelines of the University of Utah Institutional Animal Care and Use Committee.

### Zebrafish husbandry

Zebrafish were bred and maintained as described (Westerfield, 1993). Lines utilized throughout the study included *Tg(8xGli:mCH)* (Mich et al., 2014), *Tg(shh:H2A-GFP);Tg(ptch2:H2A-mCH)* (Gordon et al., 2018); *tp53^M214K^* (Berghmans et al., 2005); *Tg(atoh1c:Gal4)* (Kidwell et al., 2018)*, Tg(ptf1a:Gal4)* (Parsons et al., 2009), *Tg(dbx1a:GFP)* (Gribble et al., 2009), *Tg(ngn1:GFP)* (Blader et al., 2003), *Tg(olig2:dsRed)* (Kim et al., 2008), and *Tg(actb2:loxP-GFP-loxP-TagRFP)* (Horstick et al., 2015). The *ptch1^tj222^* line was obtained from the International Zebrafish Resource Center (http://www.zebrafish.org/home/guide.php) and has been previously described (Heisenberg et al., 1996). The *ptch2(τι11)* and *ptch2(INS+3/τι7)* lines were generated as described below.

### Method details

### Genomic DNA extraction and genotyping

Genomic DNA was extracted from adult zebrafish by fin clipping. Fins were lysed in alkaline lysis solution (25 mM NaOH, 0.2 mM EDTA) at 95°C for 2 hours and neutralized with 40 mM Tris pH 5.0. Genotyping was performed using high-melt-resolution-analysis (HRMA) as previously described (Casey et al., 2022; Dahlem et al., 2012; Xing et al., 2014; Boer et al., 2015). Briefly, HRMA was used to amplify a 103-bp and 96-bp product for *ptch1(gRNA #1)* and *ptch1(gRNA #2)*, respectively. Similarly, HRMA was used to amplify a 71-bp product for *ptch^tj222^,* a 96-bp product for *ptch2,* a 109-bp product for *grk3(gRNA #e3),* a 123-bp product for *grk3(gRNA #e12)*, and a 105-bp product for *grk3(gRNA #e18). grk3 (gRNA #e3, gRNA #e12,* and *gRNA #e18)*, *ptch1 (gRNA #1),* and *ptch1^tj222^* were cycled with the following conditions: denaturation at 96°C for 5 minutes; 55 cycles of 30 seconds at 96°C, 30 seconds at 68°C, 30 seconds at 72°C, ending with 3 minutes at 95°C and cooled to 4°C. *Ptch2* and *ptch1 (gRNA #2)* were cycled with: denaturation at 94°C for 3 minutes; 50 cycles of 30 seconds at 94°C, 20 seconds at 64°C (*ptch2*) or 70°C(*ptch1 gRNA #2*), ending with 30 seconds at 94°C, 30 seconds at 25°C and cooled to 4°C. HRMA results were validated by standard polymerase chain reaction (PCR) of genomic DNA and Sanger sequencing.

### RNA sequencing

RNA was isolated from three tumor-bearing ptch1 crispant brains and three normal zebrafish brains using the QIAGEN miRNeasy micro kit (217084). RNA quality was assessed using the Agilent RNA ScreenTape Assay (Agilent 5067-5579 and 5067-5580). The Illumina TruSeq Stranded Total RNA kit with Ribo-Zero Gold (Illumina 20020598) was used for library preparation. Sequencing libraries were chemically denatured and applied to Illumina NovaSeq flow cells using the NovaSeq XP chemistry workflow (Illumina 20021665). The flow cell was transferred to an Illumina NovaSeq instrument and a 2 x 150 cycle paired end sequence run with 100 M reads was performed using a NovaSeq S2 reagent kit (Illumina 20012860). DESeq2 was used to perform differential gene expression analysis on ptch1 tumor-bearing zebrafish compared to controls or NB-FOXR2 CNS-PNET tumors (Love et al., 2014; Modzelewska et al., 2016). A heatmap of the top 40 differentially regulated genes was generated in RStudio. Tumors were not able to be distinguished from normal brain tissue therefore, animals were deemed to be tumor-bearing due to a protrusion of the brain and abnormal swimming behavior.

To visualize the mutational spectrum in the *ptch1* tumors, raw FASTQ *ptch1-*mutant RNA sequencing files were aligned to the zebrafish reference genome to create a BAM file. BAM files were loaded into the Integrated Genomics Viewer (IGV) tool and again aligned to the appropriate version of the zebrafish reference genome (Robinson et al., 2011). We zoomed to exon 6 of the *ptch1* locus where gRNA #1 targets and visualized the insertions/deletions in each individual read in the alignment track. Insertions/deletions were tabulated for every available read of each *ptch1-*mutant sample and the number of reads identified per insertion/deletion was noted.

### GSEA

The homologene and biomaRt packages were run to convert zebrafish gene names to human gene names (Durinck et al., 2009; Sayers et al., 2022). Zebrafish gene names without a human homolog were not included in the final rank list. Duplicate genes with the highest log2FC value were retained. Rank lists were generated using ptch1 tumor versus normal brain or ptch1 tumor versus the zebrafish NB-FOXR2 CNS-PNET model driven by NRAS (Modzelewska et al., 2016). Analysis was performed using GSEA (v4.2.2).

Lists of zebrafish midbrain-hindbrain boundary (MHB) genes and hindbrain glutamatergic neuron markers were curated from the available literature. Zebrafish gene names were converted to human gene names and GSEA was run with the rank lists described above.

### PCA

To determine which human medulloblastoma subtype is modeled by the zebrafish *ptch1* tumors, PCA of human medulloblastoma and zebrafish *ptch1* crispant samples was performed based on the genes differentially expressed between *ptch1* tumors and our previously published zebrafish NRAS tumors (Modzelewska et al., 2016). Human microarray expression (Cavalli et al., 2017; GEO: GSE85217) and zebrafish RNA-seq log2 normalized counts were joined together, and only genes that exist in both human microarray and zebrafish RNA-seq data were kept. PCA data was generated using an in-house R package plot_pca (https://github.com/HuntsmanCancerInstitute/hciR).

### Immunohistochemistry

Paraffin-embedded zebrafish sections were generated and hematoxylin-and-eosin staining was performed by the Histology Core at the Huntsman Cancer Institute. Slides were imaged on a Zeiss Axio Scan.Z1 microscope at a magnification of 10X and 20X.

### Tumor Survival

To generate survival curves, fish were visually monitored in tanks weekly. Animals were sacrificed when the tumor impaired the animals ability to swim or behave normally. Statistical significance was calculated using the Mantel-Cox test with GraphPad Prism 9.

### gRNA target site design and preparation

gRNA target sites were designed using CHOPCHOP (http://chopchop.cbu.uib.no) or the IDT custom gRNA design tool as previously described (Hoshijima et al., 2019; Labun et al., 2019). Briefly, target sites were selected that had low self-complementarity, a GC content between 45-75%, no off-targets, a predicted efficiency of 45% or greater, and contained a GGG or AGG PAM sequence. Alt-R crRNA and Alt-R tracrRNA were synthesized by IDT. crRNA:tracrRNA duplexes and gRNA:Cas9 RNP complexes were prepared as previously described (Hoshijima et al., 2019).

### Microinjections

25μM gRNA:Cas9 RNP complexes were prepared as previously described (Hoshijima et al., 2019) and injected into the single cell at a dosage of 50 to 65 ng. Briefly, crRNA was annealed to tracrRNA by heating equal volumes at 95°C for 5 mins, cooling at 0.1°C/sec to 25°C, incubating for 5 mins at 25°C, and cooling to 4°C. Double gRNA injections utilized a total of 25μM-37.5μM gRNA (12.5μM-25μM gRNA each). Triple gRNA injections utilized a total of 62.5μM-75μM gRNA (12.5μM-25μM gRNA each). Prior to injection, gRNA and Cas9 were incubated at 37°C for 5 minutes for form the RNP complex.

### CRISPR/Cas9

crRNA was designed against exon 6 and exon 23 of *ptch1* (GRCz10 transcript 201) (GAGCTGATAATGGGCAGTCGGGG and TCTCGTCAAAGGGCACGTGAGGG, respectively), and against exon 16 of *ptch2* (GRCz10 transcript 201) (TTATCATGGATCCACTCCCGAGG). crRNA was designed against exon 3, exon 12 and exon 18 of *grk3* (GRCz11 transcript 202) (CTGCATGAACGAGATCGACGAGG, ACACGTCCGCATCTCTGACCTGG, and ATAAAAACGAGGCTCGCAAGAGG respectively). crRNA was designed against exon 3 of inka1b (GRCz11 transcript 201) (GGAGAATCACGCTGAACGTTTGG) (negative control) and exon 8 of smo (GRCz11 transcript 201) (CTTCTTCAATCAAGCTGAGTGGG) (positive control). crRNA was synthesized by IDT and annealed to tracrRNA (IDT) as described above.

### Founder selection for ptch2 mutations

Single adult primary injected ptch2 crispants were outcrossed and progeny were assessed for transmission of mutations via HRMA. Crosses that resulted in at least 37.5% of the tested embryos with a mutation were grown as the F1 generation. F1 heterozygous gDNA was PCR amplified followed by Sanger sequencing to determine the exact mutation. Ptch2 heterozygous F1 animals were incrossed to generate the F2 generation containing a deletion of 11 bp (Chr 2: 33975673-33975684) or a deletion of 7 bp (Chr 2: 33975679-33975685) coupled with an insertion of ATG.

### Image acquisition and processing

Brightfield images were taken using an Olympus SZX16 microscope with an aperture of 0.3 and configured with an Olympus DP74-CU camera. Fluorescent images were taken using an Olympus Fluoview FV1200 confocal microscope with a UPlanSApo 10X/0.40 or UPlanSApo 60X/1.20W objective. H&E images were taken using a Zeiss Axio Scan.Z1 slide scanner at a magnification of 10X and 20X. Figures were generated using Adobe Illustrator (2022) and Adobe Photoshop (2022).

### Statistical Analysis

The Mantel-Cox test was used to determine the statistical significance of survival data. An unpaired two-tailed t-test with Welch’s correction was used to calculate the significance of animals with pupil size differences. All statistical analysis was calculated using Prism 9.

## Supplemental figure legends

**Supplemental Figure 1, related to Figure 1 and 2. The *ptch1* gene is disrupted in *ptch1-*crispant tumors.** (A) Schematic of the mutational spectrum of the *ptch1* gene identified from *ptch1-*crispant whole brain RNA sequencing. A partial sequence of exon 6 is shown with the gRNA #1 target sequence highlighted. Identified deletions (top) and insertions (bottom) are shown along with the number of reads associated with that mutation and their predicted outcome on the Ptch1 protein (coding, green; non-coding, red). (B) Schematic of the zebrafish brain with important cerebellar structures highlighted. (C) (Left) Bright-field image of 6-to-7-wpf control animals or animals with the indicated genetic backgrounds that were injected at the one-cell stage with either *ptch1* gRNA #1 alone. (Right) Bright-field image of 6-to-7-wpf control animals or animals with the indicated genetic backgrounds that were injected at the one-cell stage with *ptch1* gRNAs #1 and #2. (Middle and right) Sagittal section of control brain or *ptch1*-crispant brain tumors stained with hematoxylin and eosin. White arrowheads indicate location of the tumor in the left panels. White boxes in the middle panel are shown at higher magnification in the right panel. Dashed black lines demarcate the cerebellum if discernible.

**Supplemental Figure 2, related to Figure 1. Transient and germline *ptch2-*mutant zebrafish do not develop tumors.**

(A) Schematic of the Ptch2 protein domains and corresponding exons. The gRNA target site is indicated. N, N-terminal domain; TM, transmembrane domain; ECD1 and ECD2, extracellular domains 1 and 2; C, C-terminal domain.

(B) Schematic of the predicted Ptch2 mutant proteins and the corresponding DNA sequencing for each of the mutant *ptch2* alleles.

(C) Chart indicating the percentage of surviving *ptch2* wild-type, heterozygous, and mutant adults identified from heterozygous incrosses of each of the indicated lines.

(D) Representative images of the coloboma eye phenotype (see arrowheads) in *ptch2*-homozygous-mutant and *ptch2-*crispant zebrafish larvae. Animals were derived from heterozygous incrosses of each of the indicated lines or from *ptch2* gRNA injected animals. Scale bar, 100μm.

**Supplemental Figure 3, related to Figure 4. Zebrafish *ptch1*-crispant tumors have characteristics of differentiated neurons.**

(A) Heatmap of the top 40 differentially regulated genes in *tp53^M214K^*; *ptch1*-crispant (gRNA #1) whole brains and *tp53^M214K^*; *NRAS^WT^* brain tumors (Modzelewska et al., 2016). Boxes highlight OPC markers (*sox10* and *erbb3)* that are upregulated in the CNS NB-FOXR2 brain tumors.

(B) GSEA was performed on genes that were differentially expressed between *tp53^M214K^; ptch1*-crispant (gRNA #1) whole brains and *tp53^M214K^; NRAS^WT^* brain tumors.

**Supplemental Figure 4, related to Figure 4. SHH-producing and -responding cells in the zebrafish hindbrain.**

(A) Lateral and (B) dorsal views of a representative 6-dpf *Tg(shh:H2A-GFP); Tg(8xGli:mCH*) zebrafish. The green fluorescent SHH-producing cells generated by the *shh:H2A-GFP* transgene are shown on the left, red fluorescent SHH-responsive cells generated by the *8xGli:mCH* transgene are shown in the middle, and finally a merged view of green and red fluorescent cells in the middle right. Lateral and dorsal views of 6-dpf *Tg(8xGli:mCH);Tg(actb2:loxP-GFP-loxP-TagRFP)* zebrafish are shown on the far right for spatial clarity of the forebrain (FB), midbrain (MB) and hindbrain (HB). Arrows indicate the location of the SHH-responsive cells shown in the left panels.

**Supplemental Figure 5, related to Figure 4. Zebrafish *ptch1-*crispant tumors are enriched for midbrain-hindbrain boundary genes.**

(A-B) (Top) GSEA identified enrichment for midbrain-hindbrain boundary genes among the differentially expressed genes between (A) control (*tp53^M214K^;Casper*) whole brains and *tp53^M214K^; ptch1-*crispant (gRNA #1) whole brains and (B) *tp53^M214K^; ptch1-*crispant (gRNA #1) whole brains versus *tp53^M214K^; NRAS^WT^* tumors.

**Supplemental Table 1. List of differentially expressed genes, related to Figure 4**. Genes were filtered to include those that were protein-coding and with *p*-adj<0.05.

**Supplemental Table 2. Hindbrain glutamatergic neurons gene list, related to Figure S3.** Genes expressed in only glutamatergic neurons in the hindbrain of 8 dpf zebrafish (Zhang et al., 2021).

**Supplemental Table 3. Midbrain hindbrain boundary gene list with accompanying references, related to Figure S5.**

**Supplemental Table 4. List of primers.**

**Supplemental Table 5. List of crRNAs.**

