## Supplemental Figures for "A Simple and Scalable Zebrafish Model of Sonic Hedgehog Medulloblastoma"

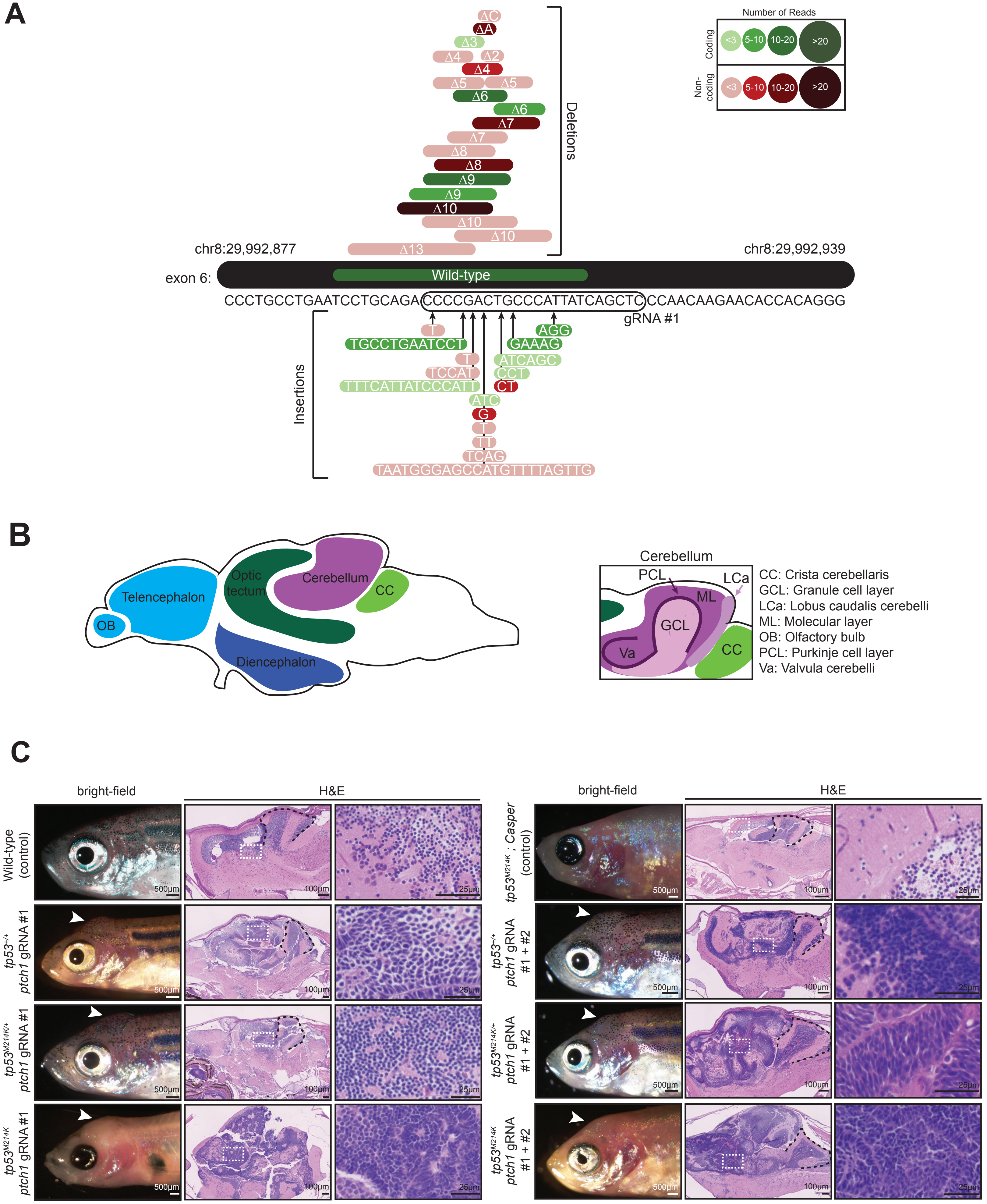
A
Number of Reads
Coding
Non-coding
<3
5-10
10-20
>20
Deletions
ΔC
ΔA
Δ3
Δ4
Δ2
Δ4
Δ5
Δ5
Δ6
Δ6
Δ7
Δ7
Δ8
Δ8
Δ9
Δ9
Δ10
Δ10
Δ10
Δ13
chr8:29,992,877
chr8:29,992,939
exon 6:
Wild-type
CCCTGCCTGAATCCTGCAGACCCCGACTGCCCATTATCAGCTCCCAACAAGAACACCACAGGG
gRNA #1
Insertions
T
AGG
TGCCTGAATCCT
GAAAG
T
ATCAGC
TCCAT
CCT
TTTCATTATCCCATT
CT
ATC
G
T
TT
TCAG
TAATGGGAGCCATGTTTTAGTTG
B
Telencephalon
OB
Optic tectum
Cerebellum
CC
Diencephalon
Cerebellum
PCL
LCa
ML
GCL
Va
CC
CC: Crista cerebellaris
GCL: Granule cell layer
LCa: Lobus caudalis cerebelli
ML: Molecular layer
OB: Olfactory bulb
PCL: Purkinje cell layer
Va: Valvula cerebelli
C
bright-field
H&E
Wild-type (control)
tp53+/+ ptch1 gRNA #1
tp53M214K/+ ptch1 gRNA #1
tp53M214K ptch1 gRNA #1
tp53M214K ; Casper (control)
tp53+/+ ptch1 gRNA #1 + #2
tp53M214K/+ ptch1 gRNA #1 + #2
tp53M214K ptch1 gRNA #1 + #2
500μm
100μm
25μm

**Supplemental Figure 1, related to Figure 1 and 2. The *ptch1* gene is disrupted in *ptch1*-crispant tumors.** (A) Schematic of the mutational spectrum of the *ptch1* gene identified from *ptch1*-crispant whole brain RNA sequencing. A partial sequence of exon 6 is shown with the gRNA #1 target sequence highlighted. Identified deletions (top) and insertions (bottom) are shown along with the number of reads associated with that mutation and their predicted outcome on the Ptch1 protein (coding, green; non-coding, red). (B) Schematic of the zebrafish brain with important cerebellar structures highlighted. (C) (Left) Bright-field image of 6-to-7-wpf control animals or animals with the indicated genetic backgrounds that were injected at the one-cell stage with either *ptch1* gRNA #1 alone. (Right) Bright-field image of 6-to-7-wpf control animals or animals with the indicated genetic backgrounds that were injected at the one-cell stage with *ptch1* gRNAs #1 and #2. (Middle and right) Sagittal section of control brain or *ptch1*-crispant brain tumors stained with hematoxylin and eosin. White arrowheads indicate location of the tumor in the left panels. White boxes in the middle panel are shown at higher magnification in the right panel. Dashed black lines demarcate the cerebellum if discernible.

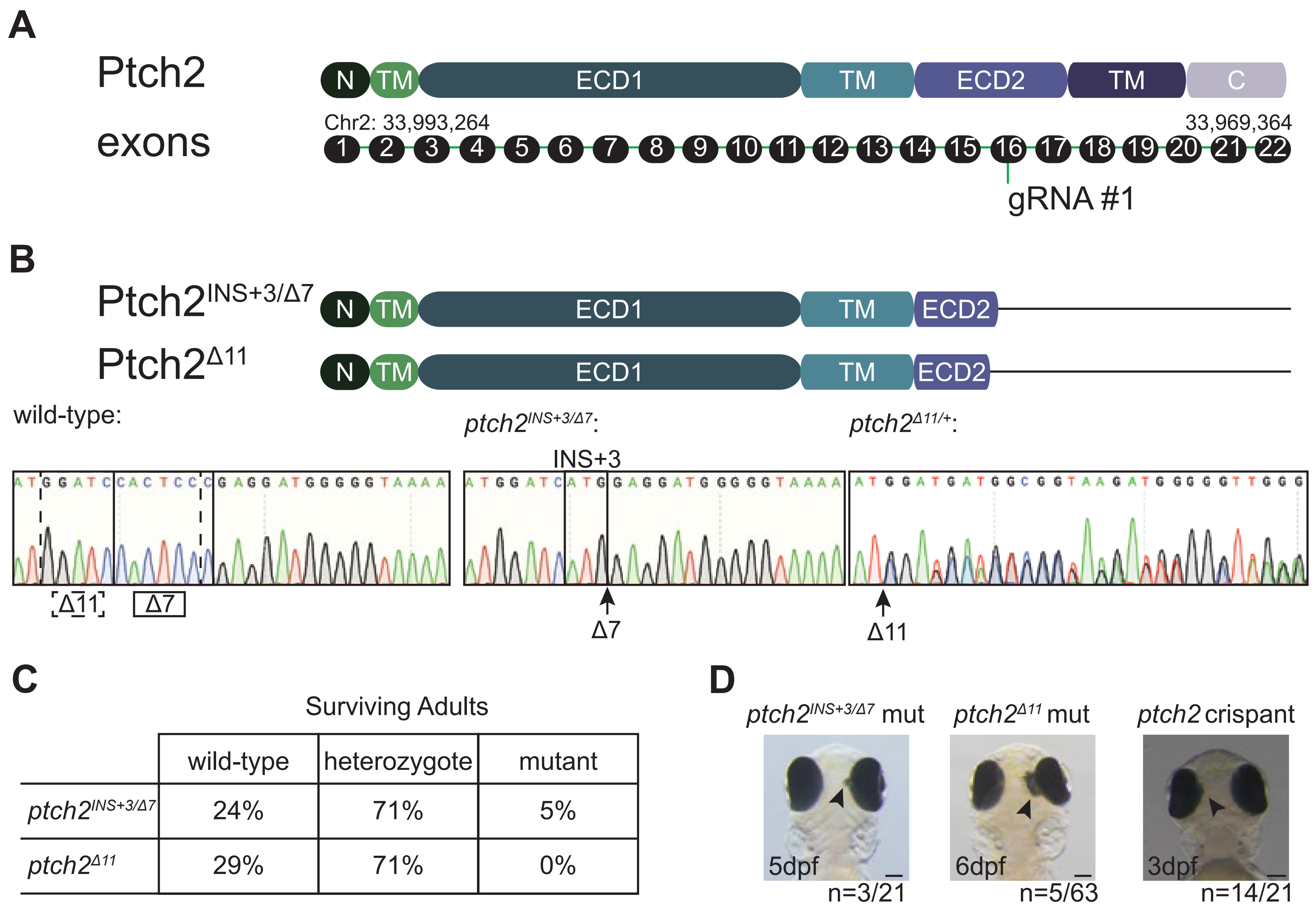


| | wild-type | heterozygote | mutant |
|---|---|---|---|
| *ptch2*$^{INS+3/\Delta7}$ | 24% | 71% | 5% |
| *ptch2*$^{\Delta11}$ | 29% | 71% | 0% |

**Supplemental Figure 2, related to Figure 1. Transient and germline *ptch2*-mutant zebrafish do not develop tumors.**

(A) Schematic of the Ptch2 protein domains and corresponding exons. The gRNA target site is indicated. N, N-terminal domain; TM, transmembrane domain; ECD1 and ECD2, extracellular domains 1 and 2; C, C-terminal domain.

(B) Schematic of the predicted Ptch2 mutant proteins and the corresponding DNA sequencing for each of the mutant *ptch2* alleles.

(C) Chart indicating the percentage of surviving *ptch2* wild-type, heterozygous, and mutant adults identified from heterozygous incrosses of each of the indicated lines.

(D) Representative images of the coloboma eye phenotype (see arrowheads) in *ptch2*-homozygous-mutant and *ptch2*-crispant zebrafish larvae. Animals were derived from heterozygous incrosses of each of the indicated lines or from *ptch2* gRNA injected animals. Scale bar, 100μm.

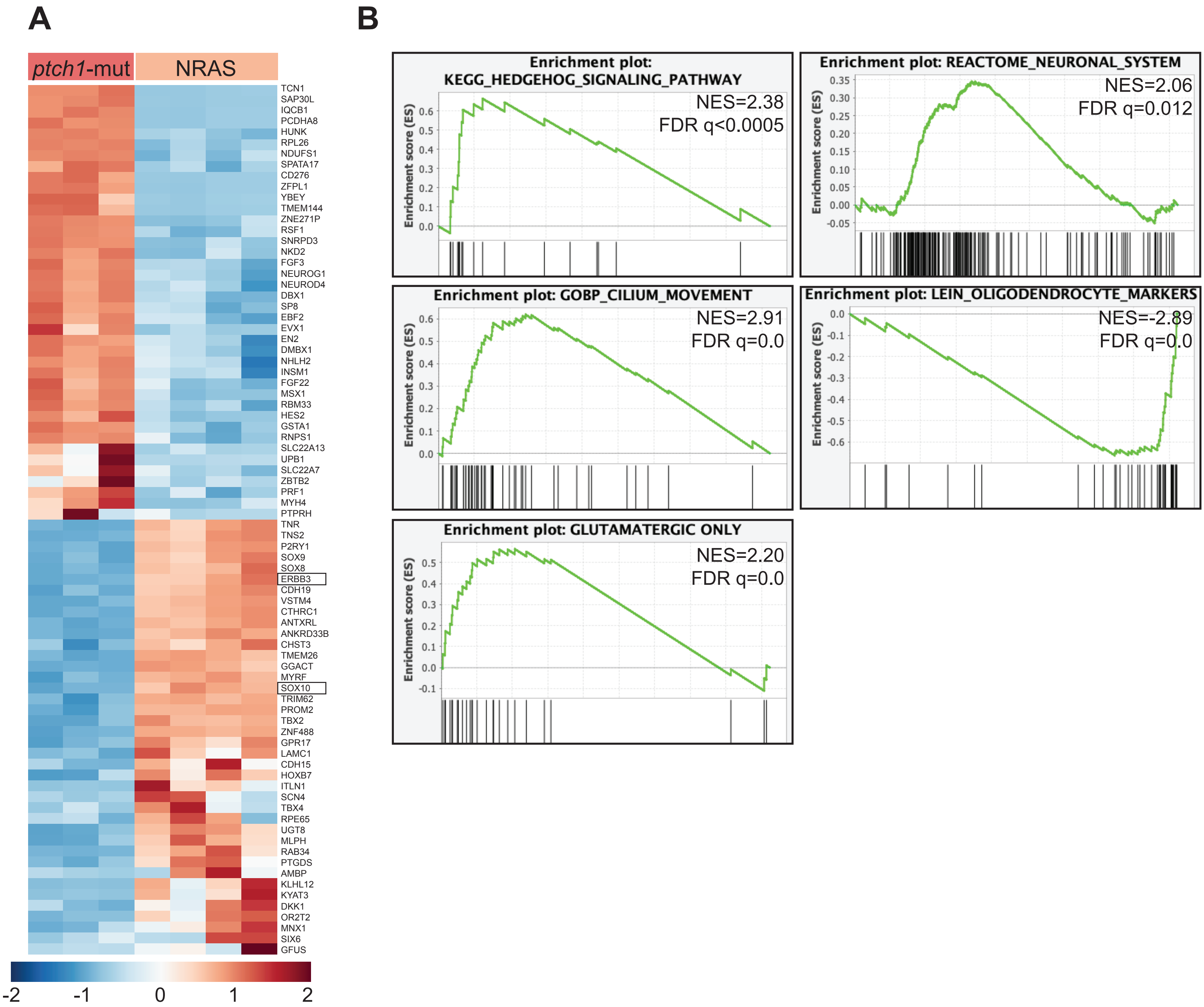

A
ptch1-mut
NRAS
TCN1
SAP30L
IQCB1
PCDHA8
HUNK
RPL26
NDUFS1
SPATA17
CD276
ZFPL1
YBEY
TMEM144
ZNE271P
RSF1
SNRPD3
NKD2
FGF3
NEUROG1
NEUROD4
DBX1
SP8
EBF2
EVX1
EN2
DMBX1
NHLH2
INSM1
FGF22
MSX1
RBM33
HES2
GSTA1
RNPS1
SLC22A13
UPB1
SLC22A7
ZBTB2
PRF1
MYH4
PTPRH
TNR
TNS2
P2RY1
SOX9
SOX8
ERBB3
CDH19
VSTM4
CTHRC1
ANTXRL
ANKRD33B
CHST3
TMEM26
GGACT
MYRF
SOX10
TRIM62
PROM2
TBX2
ZNF488
GPR17
LAMC1
CDH15
HOXB7
ITLN1
SCN4
TBX4
RPE65
UGT8
MLPH
RAB34
PTGDS
AMBP
KLHL12
KYAT3
DKK1
OR2T2
MNX1
SIX6
GFUS
-2
-1
0
1
2
B
Enrichment plot: KEGG_HEDGEHOG_SIGNALING_PATHWAY
NES=2.38
FDR q<0.0005
Enrichment plot: REACTOME_NEURONAL_SYSTEM
NES=2.06
FDR q=0.012
Enrichment plot: GOBP_CILIUM_MOVEMENT
NES=2.91
FDR q=0.0
Enrichment plot: LEIN_OLIGODENDROCYTE_MARKERS
NES=-2.89
FDR q=0.0
Enrichment plot: GLUTAMATERGIC ONLY
NES=2.20
FDR q=0.0
Enrichment score (ES)

**Supplemental Figure 3, related to Figure 4. Zebrafish *ptch1*-crispant tumors have characteristics of differentiated neurons.**

(A) Heatmap of the top 40 differentially regulated genes in *tp53*$^{M214K}$; *ptch1*-crispant (gRNA #1) whole brains and *tp53*$^{M214K}$; *NRAS*$^{WT}$ brain tumors (Modzelewska et al., 2016). Boxes highlight OPC markers (*sox10* and *erbb3)* that are upregulated in the CNS NB-FOXR2 brain tumors.

(B) GSEA was performed on genes that were differentially expressed between *tp53*$^{M214K}$*; ptch1*-crispant (gRNA #1) whole brains and *tp53*$^{M214K}$*; NRAS*$^{WT}$ brain tumors.

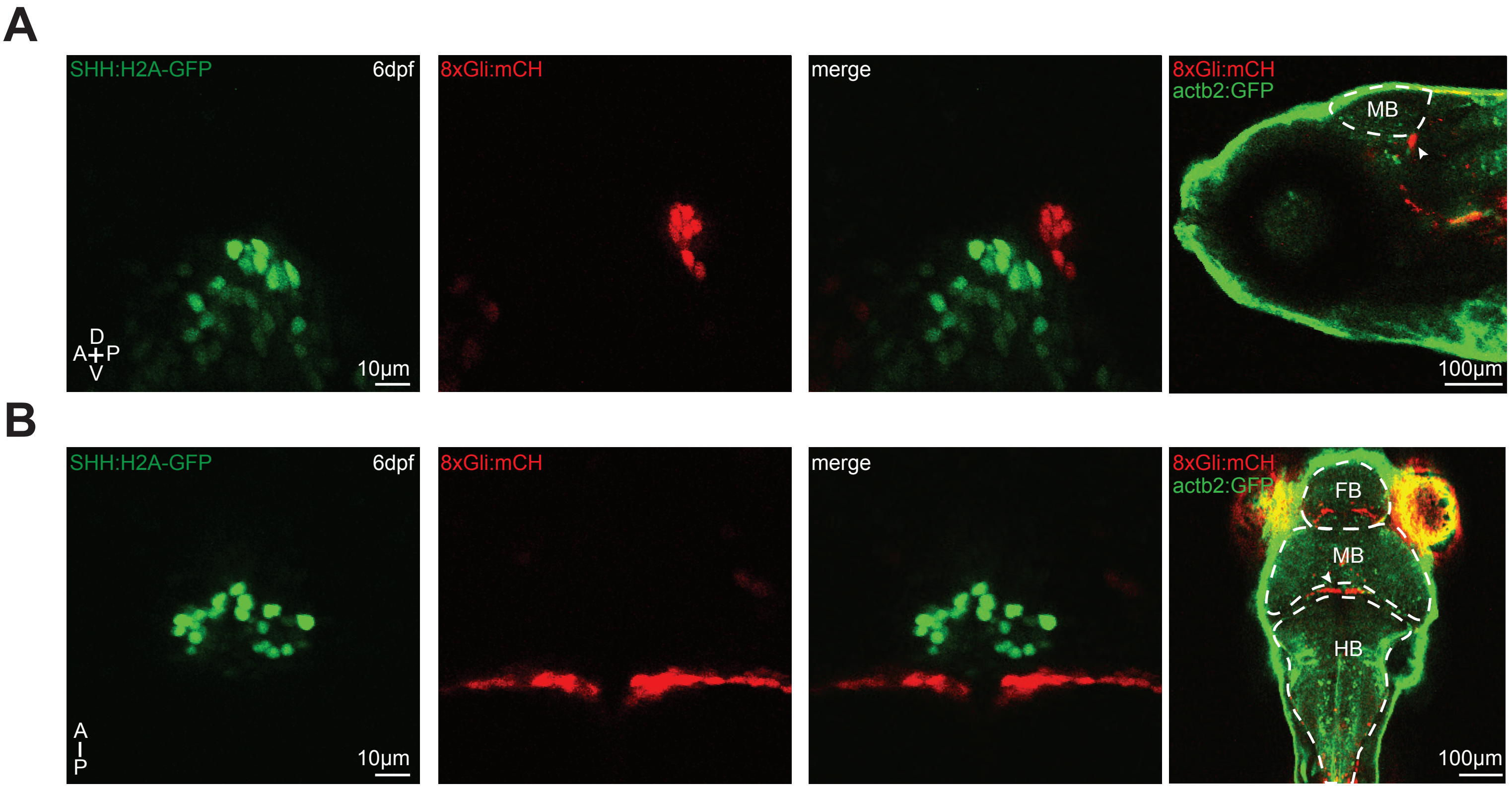

A
SHH:H2A-GFP
6dpf
8xGli:mCH
merge
8xGli:mCH
actb2:GFP
MB
D
A
P
V
10μm
100μm
B
SHH:H2A-GFP
6dpf
8xGli:mCH
merge
8xGli:mCH
actb2:GFP
FB
MB
HB
A
P
10μm
100μm

**Supplemental Figure 4, related to Figure 4. SHH-producing and -responding cells in the zebrafish hindbrain.**

(A) Lateral and (B) dorsal views of a representative 6-dpf *Tg(shh:H2A-GFP); Tg(8xGli:mCH*) zebrafish. The green fluorescent SHH-producing cells generated by the *shh:H2A-GFP* transgene are shown on the left, red fluorescent SHH-responsive cells generated by the *8xGli:mCH* transgene are shown in the middle, and finally a merged view of green and red fluorescent cells in the middle right. Lateral and dorsal views of 6-dpf *Tg(8xGli:mCH);Tg(actb2:loxP-GFP-loxP-TagRFP)* zebrafish are shown on the far right for spatial clarity of the forebrain (FB), midbrain (MB) and hindbrain (HB). Arrows indicate the location of the SHH-responsive cells shown in the left panels.

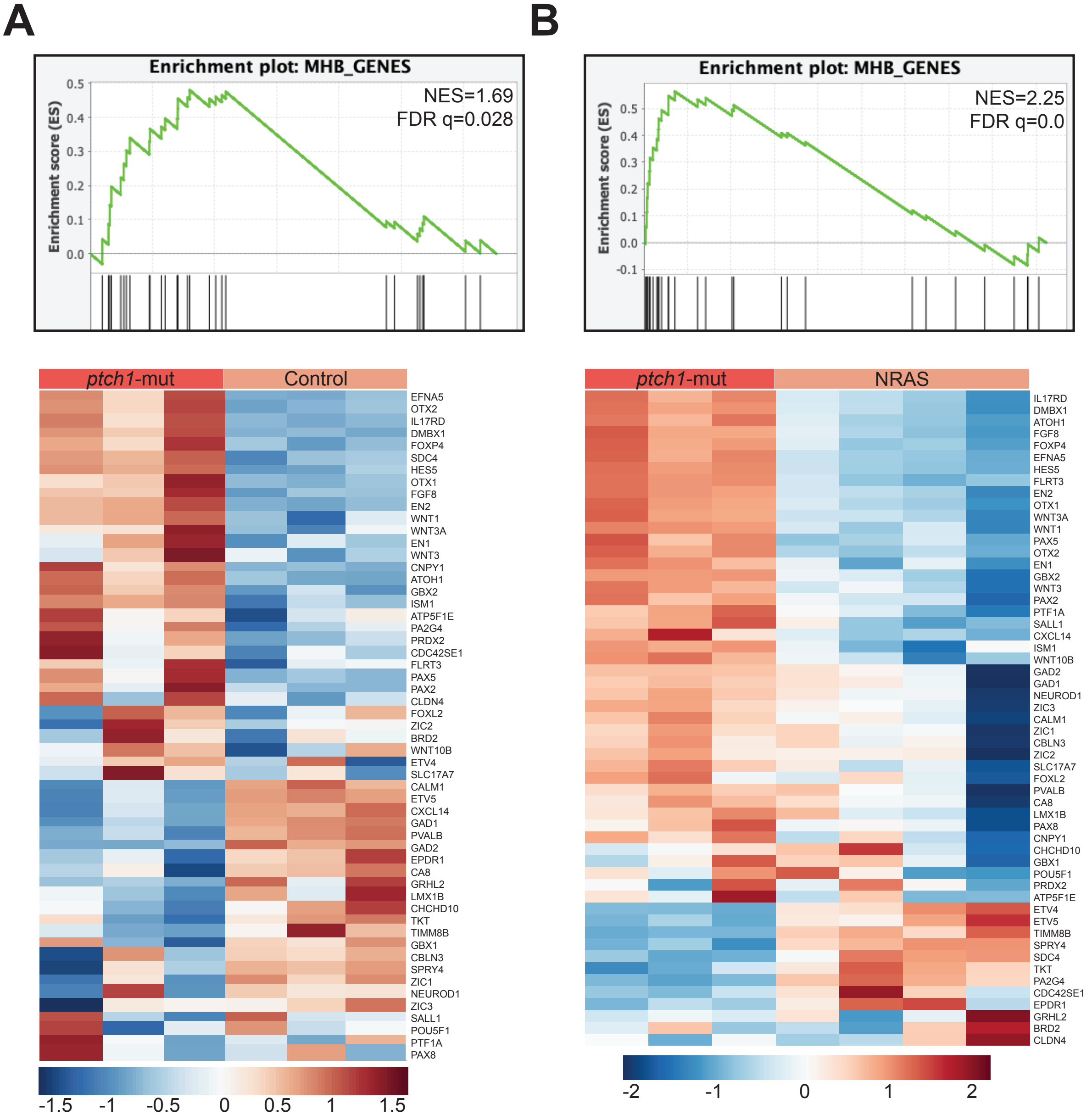
A
Enrichment plot: MHB_GENES
Enrichment score (ES)
NES=1.69
FDR q=0.028
ptch1-mut
Control
EFNA5
OTX2
IL17RD
DMBX1
FOXP4
SDC4
HES5
OTX1
FGF8
EN2
WNT1
WNT3A
EN1
WNT3
CNPY1
ATOH1
GBX2
ISM1
ATP5F1E
PA2G4
PRDX2
CDC42SE1
FLRT3
PAX5
PAX2
CLDN4
FOXL2
ZIC2
BRD2
WNT10B
ETV4
SLC17A7
CALM1
ETV5
CXCL14
GAD1
PVALB
GAD2
EPDR1
CA8
GRHL2
LMX1B
CHCHD10
TKT
TIMM8B
GBX1
CBLN3
SPRY4
ZIC1
NEUROD1
ZIC3
SALL1
POU5F1
PTF1A
PAX8
-1.5 -1 -0.5 0 0.5 1 1.5
B
Enrichment plot: MHB_GENES
Enrichment score (ES)
NES=2.25
FDR q=0.0
ptch1-mut
NRAS
IL17RD
DMBX1
ATOH1
FGF8
FOXP4
EFNA5
HES5
FLRT3
EN2
OTX1
WNT3A
WNT1
PAX5
OTX2
EN1
GBX2
WNT3
PAX2
PTF1A
SALL1
CXCL14
ISM1
WNT10B
GAD2
GAD1
NEUROD1
ZIC3
CALM1
ZIC1
CBLN3
ZIC2
SLC17A7
FOXL2
PVALB
CA8
LMX1B
PAX8
CNPY1
CHCHD10
GBX1
POU5F1
PRDX2
ATP5F1E
ETV4
ETV5
TIMM8B
SPRY4
SDC4
TKT
PA2G4
CDC42SE1
EPDR1
GRHL2
BRD2
CLDN4
-2 -1 0 1 2

**Supplemental Figure 5, related to Figure 4. Zebrafish *ptch1*-crispant tumors are enriched for midbrain-hindbrain boundary genes.**

(A-B) (Top) GSEA identified enrichment for midbrain-hindbrain boundary genes among the differentially expressed genes between (A) control (*tp53*$^{M214K}$*;Casper*) whole brains and *tp53*$^{M214K}$*; ptch1*-crispant (gRNA #1) whole brains and (B) *tp53*$^{M214K}$*; ptch1*-crispant (gRNA #1) whole brains versus *tp53*$^{M214K}$*; NRAS*$^{WT}$ tumors.

**Supplemental Table 2. Hindbrain glutamatergic neurons gene list, related to Figure S3.** Genes expressed in only glutamatergic neurons in the hindbrain of 8 dpf zebrafish (Zhang et al., 2021).

| Human homologs |
| --- |
| ACLY |
| ADCYAP1 |
| ARHGEF1 |
| ATP1B1 |
| BDNF |
| CALB2 |
| CALM2 |
| CAMK2N1 |
| CBLN1 |
| CBLN2 |
| DDIT3 |
| DRGX |
| EGR2 |
| ESRRB |
| EYA1 |
| FNDC4 |
| FOXD3 |
| FOXP4 |
| FSTL5 |
| GBX2 |
| GRM5 |
| HMX2 |
| HPCAL4 |
| HUNK |
| IRX1 |
| IRX2 |
| IRX4 |
| KCTD12 |
| LMO3 |
| LMX1B |
| MAFB |
| MYCN |
| NANOS1 |
| NR2F1 |
| NRGN |
| NRN1 |
| NTNG1 |
| OLFM3 |
| PIK3R3 |
| PITX2 |
| PKIB |

| |
|---|
| POU4F1 |
| PPP1R1C |
| PTGER4 |
| QDPR |
| R3HDM1 |
| RARA |
| RORA |
| RPRM |
| SHOX2 |
| SLC17A6 |
| SOX4 |
| TFAP2A |
| TFAP2B |
| TLX2 |
| TLX3 |
| VAV3 |

**Supplemental Table 3. Midbrain hindbrain boundary gene list with accompanying references, related to Figure S5.**

| Human Homologs | Reference |
|---|---|
| EN2 | Tambalo et al., 2020; Belting et al., 2001 |
| IL17RD | Tambalo et al., 2020; Gibbs et al., 2017 |
| PAX5 | Tambalo et al., 2020; Brand et al., 1996 |
| PAX8 | Tambalo et al., 2020; O’Hara et al., 2005 |
| OTX1 | Tambalo et al., 2020; Rhinn & Brand 2001 |
| CNPY1 | Tambalo et al., 2020; Hirate et al., 2006 |
| PAX2 | Tambalo et al., 2020; Belting et al., 2001; Brand et al., 1996 |
| FGF8 | Tambalo et al., 2020; Belting et al., 2001; O’Hara et al., 2005 |
| GBX2 | Tambalo et al., 2020; Rhinn & Brand 2001 |
| FLRT3 | Tambalo et al., 2020 |
| ISM1 | Tambalo et al., 2020 |
| CHCHD10 | Tambalo et al., 2020 |
| EFNA5 | Tambalo et al., 2020 |
| SDC4 | Tambalo et al., 2020 |
| CLDN4 | Tambalo et al., 2020 |
| CALM1 | Tambalo et al., 2020 |
| PRDX2 | Tambalo et al., 2020 |
| CXCL14 | Tambalo et al., 2020 |
| ATP5F1E | Tambalo et al., 2020 |
| TKT | Tambalo et al., 2020 |
| PA2G4 | Tambalo et al., 2020 |
| TIMM8B | Tambalo et al., 2020 |
| FOXP4 | Tambalo et al., 2020 |
| EN1 | Tambalo et al., 2020; Brand et al., 1996 |
| POU5F1 | Belting et al., 2001 |
| HES7 | Geling et al., 2003; Geling et al., 2004; Ninkovic et al., 2005 |
| WNT1 | Belting et al., 2001; Brand et al., 1996 |
| HES5 | Sigloch et al., 2023 |
| PTF1A | Kaslin et al., 2013 |
| OTX2 | Belting et al., 2001; Rhinn & Brand 2001 |
| GBX1 | Belting et al., 2001; Rhinn & Brand 2001 |
| FOXL2 | Zhou et al., 2022 |
| WNT3 | O’Hara et al., 2005; Buckles et al. 2004 |
| WNT3A | O’Hara et al., 2005; Buckles et al. 2004 |
| WNT10B | O’Hara et al., 2005; Buckles et al. 2004 |
| GRHL2 | Dworkin et al. 2012 |
| CDC42SE1 | Dworkin et al. 2012 |
| BRD2 | Murphy et al. 2017 |
| LMX1B | O’Hara et al., 2005; Harada et al., 2016 |
| ETV4 | Harada et al., 2016 |
| ATOH1 | Kidwell et al., 2018 |

| | |
|---|---|
| NEUROD1 | Feng et al., 2022 |
| ZIC1 | Drummond et al., 2013 |
| ZIC2 | Drummond et al., 2013 |
| ZIC3 | Drummond et al., 2013 |
| EPDR1 | Ota et al., 2016 |
| PVALB | Kyostila et al., 2015 |
| CBLN3 | Dohaku et al., 2019 |
| SLC17A7 | Kyostila et al., 2015 |
| GAD2 | Fernandes et al., 2013 |
| GAD1 | Fernandes et al., 2013 |
| CA8 | Sassen et al., 2017 |
| ETV5 | Jaszai et al., 2003 |
| SALL1 | Jaszai et al., 2003 |
| SPRY4 | Jaszai et al., 2003 |
| DMBX1 | Jaszai et al., 2003 |

**Supplemental Table 4. List of primers**

| Primer Name | Sequence | Use |
|---|---|---|
| MS_ptch1_F2 | GTCCACAATGACCCACACTTCT | HRMA of ptch1 gRNA#2 |
| MS_ptch1_R2 | TTCTAGATCCTGCAGCTCCATG | HRMA of ptch1 gRNA#2 |
| Ptch1_e6_HRMAF1 | GCAGATGTGGGTCAAGGTTACA | HRMA of ptch1 gRNA#1 |
| Ptch1_e6_HRMAR2 | CTAAGCTCACCCCTGTGGTGTT | HRMA of ptch1 gRNA#1 |
| MS_ptch2_F2 | AGTAATGATCCACTGGGCTATGC | HRMA of ptch2 |
| MS_ptch2_R2 | GTTCTCTCCAGTGGTGTCGTAC | HRMA of ptch2 |
| Ptch1_seqF2 | GTCCCACAACCCCTACAACC | PCR amplification of ptch1 gRNA #2 region |
| Ptch1_seqR2 | AACGATTAACCATTCGCGTACA | PCR amplification of ptch1 gRNA #2 region |
| Ptch2_seqF2 | GATGGAGCACTGGCCTACAAG | PCR amplification of ptch2 |
| Ptch2_seqR1 | AAGGTTCTCTCCAGTGGTGTCG | PCR amplification of ptch2 |
| tj222_hrma_f1 | CATCCTCAGCATGGACCTG | HRMA of ptch1 tj222 |
| tj222_hrma_r1 | ATTACCTGACAAAGCAGCAGAA | HRMA of ptch1 tj222 |
| grk3_e12_HRMA_F2 | GGAGCGTGTGTCCTGATCTC | PCR of grk3 e12 |
| grk3_e12_HRMA_R2 | CCCCAAACAGCAGACGAATA | PCR of grk3 e12 |
| Grk3e3_hrma_fwd2 | CTCCAGGTTTCCTCCTCTTCAA | HRMA of grk3 e3 |
| Grk3e3_hrma_rev2 | GAATGCTAGCAAAGCAAAACCA | HRMA of grk3 e3 |
| Grk3_e12_hrma_F1 | CAGCCTGCGAATATCCTCTT | HRMA of grk3 e12 |
| Grk3_e12_hrma_R1 | GAGGAGGACTTGAGGAGGAAC | HRMA of grk3 e12 |
| Grk3e3_seq_fwd | CTTGCATTGTGCCTGCTTTA | PCR of grk3 e3 |
| Grk3e3_seq_rev | GTGTGGGCCATCTCATTACA | PCR of grk3 e3 |
| Grk3e18_hrma_f1 | CGAGACCGTGTATGAAGCTGTG | HRMA of grk3 e18 |
| Grk3e18_hrma_r1 | AAGTGCGGTGTCTGACCTTCTT | HRMA of grk3 e18 |
| MS_inka1b_1_F1 | CCTACGATTCTGCCTGCTGTCT | HRMA of inka1b |
| MS_inka1b_1_R1 | CAAAGTCCAGGCTCTTGATGCTG | HRMA of inka1b |

| | | |
|---|---|---|
| Inka1b_seq_F1 | CGATCAATCCAAATGCTCCT | PCR of inka1b |
| Inka1b_seq_R1 | ACTCGCCTCAAATGATGTCC | PCR of inka1b |

**Supplemental Table 5. List of crRNAs**

| Crispr identifier | Sequence | Exon Target |
|---|---|---|
| Ptch1 gRNA #2 | TCTCGTCAAAGGGCACGTGAGGG | 23 |
| Ptch1 gRNA #1 | GAGCTGATAATGGGCAGTCGGGG | 6 |
| Ptch2 gRNA #1 | TTATCATGGATCCACTCCCGAGG | 16 |
| Grk3 gRNA #2 | ACACGTCCGCATCTCTGACCTGG | 12 |
| Grk3 gRNA #3 | ATAAAAACGAGGCTCGCAAGAGG | 18 |
| Grk3 gRNA #1 | CTGCATGAACGAGATCGACGAGG | 3 |
| Inka1b gRNA #1 (Control) | GGAGAATCACGCTGAACGTTTGG | 3 |
| Smo gRNA #1 | CTTCTTCAATCAAGCTGAGTGGG | 8 |

PAM sequence is underlined
